# Unravelling hidden trophic interactions among sea urchin juveniles and macroinvertebrates by DNA amplification

**DOI:** 10.1101/2025.07.10.664104

**Authors:** Alberto Sutera, Chiara Bonaviri, Francesco Di Trapani, Francesca La Bella, Vesna Macic, Olivera Marković, Bernat Hereu, Roberto De Michele

## Abstract

Rocky reefs may shift between two distinct stable states: productive algal forests, characterized by high abundance and biodiversity of macrofauna, and impoverished barrens, dominated by overgrazing sea urchins. Barren states may persist despite the recovery of adult sea urchin predators, suggesting additional stabilizing mechanisms. Sea urchin settlers rapidly disappear in forests, but they persist in barrens, suggesting that post-settlement predation might play a crucial role in determining sea urchin population density. Visual assessment of predation events in the field is unfeasible due to the microscopic scale of both predators and preys and the complexity of the arena. In this study, we designed and tested specific primers for the detection of mtDNA of settlers of the Mediterranean sympatric sea urchin species *Paracentrotus lividus* and *Arbacia lixula* on degraded samples. By testing 360 invertebrates collected in algal forests during an urchin settling event at five mtDNA loci, we identified 60 (17%) samples positive to *P. lividus* DNA. Presence of urchin DNA was confirmed by sequencing and NGS metabarcoding analyses. Our results suggest that micropredation may represent an important process in controlling sea urchin population density and maintaining the forest state in temperate rocky reefs.

**Statements:** The data that supports the main findings of this study are available in the supplementary material of this article. Other data are available from the corresponding author upon reasonable request.

## INTRODUCTION

On rocky reefs, one of the most striking transformations is the shift from lush macroalgal forest to barren ground caused by sea urchin overgrazing (Ling et al., 2015). Forests, characterized by erect algae, boast complex architecture and high species diversity, whereas barrens are structurally simple, with low diversity, and dominated by sea urchins and encrusting organisms (Graham, 2004; Pinna et al., 2020; Steneck et al., 2002; Taylor, 1998). Transitions between these states occur globally due to various factors such as the loss of top-down control over sea urchins, habitat destruction, increases in seawater temperatures and heat waves, and the consequent changes in species physiology and distribution (Bonaviri et al., 2017; Jackson et al., 2001; Mann, 1982; Sala et al., 1998; Steneck et al., 2002). Once an ecosystem shifts, hysteresis mechanisms maintain the new state, even if initial conditions are restored (Baskett & Salomon, 2010; Scheffer & Carpenter, 2003; Suding et al., 2004). Upon establishment, barren states often see high sea urchin density and biomass, further stabilizing the state (Bonaviri et al., 2011; Gianguzza et al., 2010; Knowlton, 2004; Ling et al., 2009; Steneck et al., 2002). Research indicates that large predators of adult sea urchins can reverse urchins barrens and sustain forests (Bernstein et al., 1981; Clemente et al., 2009; Guidetti & Sala, 2007; Jackson et al., 2001; Ling et al., 2009; Mann, 1982; Sala et al., 1998; Shears & Babcock, 2002, 2003). However, despite the recovery of adult sea urchin predators, high sea urchin densities and barren communities can persist for years (Babcock et al., 2010; Foster, 1990; Pinnegar et al., 2000), suggesting the presence of other stabilizing mechanisms (Agnetta et al., 2024; Ling et al., 2015; Sala, 2004). As a matter of fact, although the role of sea urchins on structuring rocky reef communities is well known, drivers of sea urchins’ demographics are poorly understood. Recruitment and post-settlement survival are fundamental processes in sea urchin population dynamics (Hereu et al., 2012). Since urchins recruits are particularly vulnerable to predation, their mortality may represent one of the most important driver determining urchin population density (Andrew & Choat, n.d.; Harrold et al., 1991; Jennings & Hunt, 2010; Prog et al., 1987; Rowley, 1989).After the planktonic larval stage, settlement induction and recruits density appear independent of community state or conspecific abundance, with young urchin settlers equally present in both forest and barren (Balch & Scheibling, 2001; Cameron & Schroeter, 1980; Hereu et al., 2004; Hernández et al., 2010; McNaught Douglas Colin, 1999; Prado et al., 2009; Privitera et al., 2011; Rowley, 1989; Schroeter et al., 1996). However, settler population rapidly drops in forests, despite the presence of structural refuges in forests protects small urchins from predation by fishes (Boada et al., 2018; Hereu et al., 2004; Steneck et al., 2013). As a result, adult sea urchins are typically rare in forests, and abundant in barrens. It has long been postulated that invertebrate micropredators are responsible for the control of urchin settlers (Bonaviri et al., 2012; Clemente et al., 2013; Hereu et al., 2005; Jennings & Hunt, 2010; McNaught Douglas Colin, 1999; Scheibling & Hamm, 1991; Scheibling & Robinson, 2008; Steneck et al., 2013). Predation is a significant cause of early benthic invertebrate mortality (Gosselin & Qian, 1997; Griffiths & Gosselin, 2008; Hunt & Scheibling, 1997; Osman & Whitlatch, 1992, 1995, 2004; Paine, 1992; Sala & Graham, 2002). Forests, due to the greater habitat complexity, host a larger abundance and diversity of microfauna, hence potentially micropredators, than structurally simple barren grounds (Christie et al., 2009; Hauser et al., 2006; Norderhaug et al., 2012; Pinna et al.2020). Consequently, the disappearance of urchin settlers in forests might be due to predation by invertebrates (Steneck et al., 2013).

Experiments in aquaria showed that several decapod species are able to feed on urchin settlers, especially larger specimens (Bonaviri et al., 2012; Fagerli et al., 2014; Feehan et al., 2014). However, the number and variety of invertebrates tested is limited, and aquaria are artificial, oversimplified systems which poorly encompass the real trophic dynamics in the field. Traditional techniques for in situ identification of predation events involving marine invertebrates face paramount challenges, due to the small scale of both predators and preys, presence of algal canopy, frequent nocturnal activity and disturbance by the observer. Mounting of cameras on the seabed may ease some of these problems, but it still presents a limited temporal and spatial arena, often deprived of canopy and other organisms, or where preys are tethered to the substratum. As a result, predation rates measured by visual scoring might be overestimated (Fagerli et al., 2014). Analysis of gut content is problematic as well, since many invertebrates are fluid feeders, avoid consuming indigestible remains, or fully digest samples leaving no recognizable material.

In the last decades, several studies have employed environmental DNA (eDNA) to study the trophic interactions in invertebrates (Clare, 2014; Cuff et al., 2022; Pompanon et al., 2012; Symondson, 2002). Amplification of prey’s DNA by polymerase chain reaction (PCR) allows the detection of trace amounts of undigested prey material in predator guts content and in feces.

Here, we applied molecular techniques to untangle the trophic interactions involving urchins settlers and their potential micropredators. We considered the sea urchins *Paracentrotus lividus* and *Arbacia lixula*, which cause the shift from algal forest to the barren grounds in Mediterranean rocky reefs (Agnetta et al., 2015; Bonaviri et al., 2011; Hereu et al., 2008).

In particular, we aimed to: (1) design specific primers for the detection of DNA of the Mediterranean urchin species *P. lividus* and *A. lixula* in degraded samples; (2) identify the invertebrates belonging to different taxa collected during the urchin settlement stage, by both visual assignment and molecular barcoding; (3) detect potential consumers of urchin settlers.

The extent of micropredation as a process controlling urchin populations in temperate reefs is discussed.

## Materials and methods

### Sample collection

Weekly surveys by SCUBA diving along the coastline of the Montgrí massif (42.8160 N, 03.8130 E), Spain, northwestern Mediterranean Sea, and at one site on the Thyrrenian coast of Sicily (38.1086 N, 13.5382 E), Italy, central Mediterranean Sea, were conducted since the beginning of summer 2017 to detect the peak moment of sea urchin recruits abundance. In July, we observed a large number of sea urchin recruits on algal assemblages at 5–8 m depth. Invertebrates were collected by scraping an algae-covered rocky substrate by SCUBA diving, then sorted under a stereomicroscope in the laboratory on the same day of collection. Specimens were individually stored in plastic tubes in 1-10 ml of 70% ethanol, depending on the animal size, and kept at -20 °C until use. We paid attention to avoid cross-contamination between samples, by changing gloves and cleaning tweezers for each specimen.

Sea urchin juveniles were also collected in the same locations. In order to collect the small juveniles, the scraped material was placed in a salver containing a thin layer of seawater and covered with a plastic grid. After a few hours, the small sea urchins climbed actively out of the salver onto the surface of the plastic grid, probably in response to oxygen shortage (Bonaviri et al., 2012). Sea urchins were then carefully collected with tweezers and individually stored as described above. As a precaution to avoid DNA contamination with micropredators, the tubes containing sea urchins were stored separately in a different box, in a different freezer.

Adults of *P. lividus* and *A. lixula*, whose DNA was used as positive controls in primer tests, were collected at the Sicilian locality.

All animals collected in this study were invertebrates and non-cephalopods, therefore not subject to Directive 2010/63/EU restrictions. None of them belonged to endangered or protected species.

### Taxonomic identification

Invertebrates were visually classified in two rounds: at the moment of collection, and before DNA extraction. Samples were individually visualized under a stereomicroscope in the laboratory and photographed. With the aid of manuals, we identified the specimens at the lowest possible taxonomic level (Supplemental Table S1).

For molecular taxonomic identification of both urchin juveniles and potential predators by DNA barcoding, we used the “universal” *cytochrome oxidase subunit 1* (*COI1*) degenerate primer pair jgLCO1490/jgHCO2198 (Table1) described by (Geller et al., 2013), particularly effective for marine invertebrates.

### DNA extraction

For DNA extraction from both the potential predators and sea urchins, we used the Tissue DNA Kit by Ezna (VWR). Samples were removed from the ethanol bath, weighted and rinsed in 1-10 mL distilled water, depending on the animal size. For animals weighing more than 100 mg, we chopped the sample in several pieces by using a disposable blade, and processed the parts independently. Legs and claws were also removed from larger animals, whenever possible, in order to minimize the amount of host DNA recovered. Each sample was put in 2 mL plastic tube with 3 tungsten beads, and homogenized by shaking 15 s at 300 rpm at room temperature in the TissuLyser machine (Qiagen). Immediately after shaking, we added TL buffer and proceeded according to the kit protocol for tissue. DNA was resuspended in 10 mM Tris buffer, pH 9.0. DNA concentration and extraction quality were measured at a fluorometer (Synergy H1, Biotek). DNA integrity was checked by electrophoretic run in a 1% agarose gel.

### Primer design

For molecular identification of potential feeders of sea urchins, we designed several primer pairs spanning different regions of the mitochondrial gene cytochrome oxidase subunit 1 (*COI1*), commonly used in similar studies since it is highly polymorphic among different species. Due to the high polymorphism of the gene, we were concerned that different urchin accessions might also show sequence variations. In order to design primers that would work for any *P. lividus* and *A. lixula* sequence, we aligned by ClustalW (BioEdit) 254 *COI1* sequences available on NCBI, including the complete mitochondrial genome for *P. lividus* (accessions 49036145-49036397; 1028338044- 1028338084; 365735528-365735742; 564282614; 564282618; J04815.1) and 328 *COI1* sequences available for *A. lixula* (JQ745096.1-JQ745256.1; JN603630.1-JN603633.1; AF030998.1- AF031011.1; JF772935.1-JF773074.1; KU172486.1-KU172488.1; HE800533.1-HE800538.1). In the alignment, we also included the *COI1* sequences of marine invertebrate representatives of the major taxonomic group found in this survey: *Alpheus dentipes* (AF309893.1); *Amphipholis squamata* (NSECH002-13); *Ampithoe ramondi* (GBCM8446-17); *Dorvillea sp* (BAMPOL0439); *Galathea intermedia* (BNSC183-10); *Nereis pelagica*; *Ophiothrix fragilis* (LOBO010-12); *Pagurus prideaux* (JSDUK158); *Palola cf. siciliensis* (USNM1120744); *Pilumnus villosissimus* (GBCMD18798-19); *Platynereis dumerilii* (GBAN12514-19); *Synalpheus gambarelloides* (GBCDA161-12); and *Thelepus cincinnatus* (HUNTSPOL0372), in order to avoid conserved regions during primer design. Likewise, we designed primers for the mitochondrial loci *16S* and cytochrome b (*CYTb*). We designed nine different primer pairs for *P. lividus* and seven for *A. lixula*, spanning different regions of each locus and amplifying bands ranging from 63 to 178 bp in size. We intentionally selected small amplification sizes in order to maximize chances of recovery of digested and fragmented prey DNA in the guts(King et al., 2008). Details of the primers are given in Table 1.

**Table 1.**
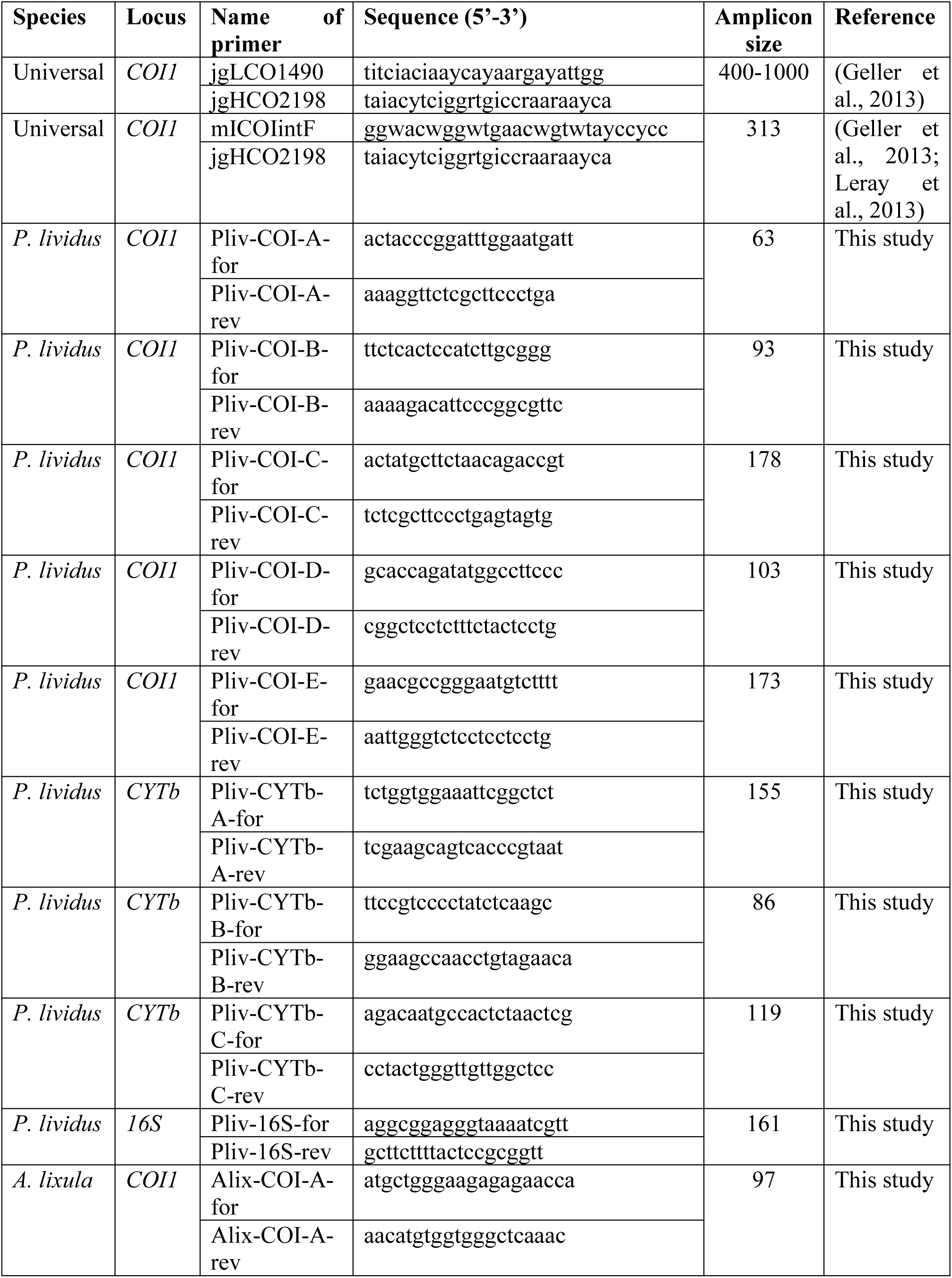
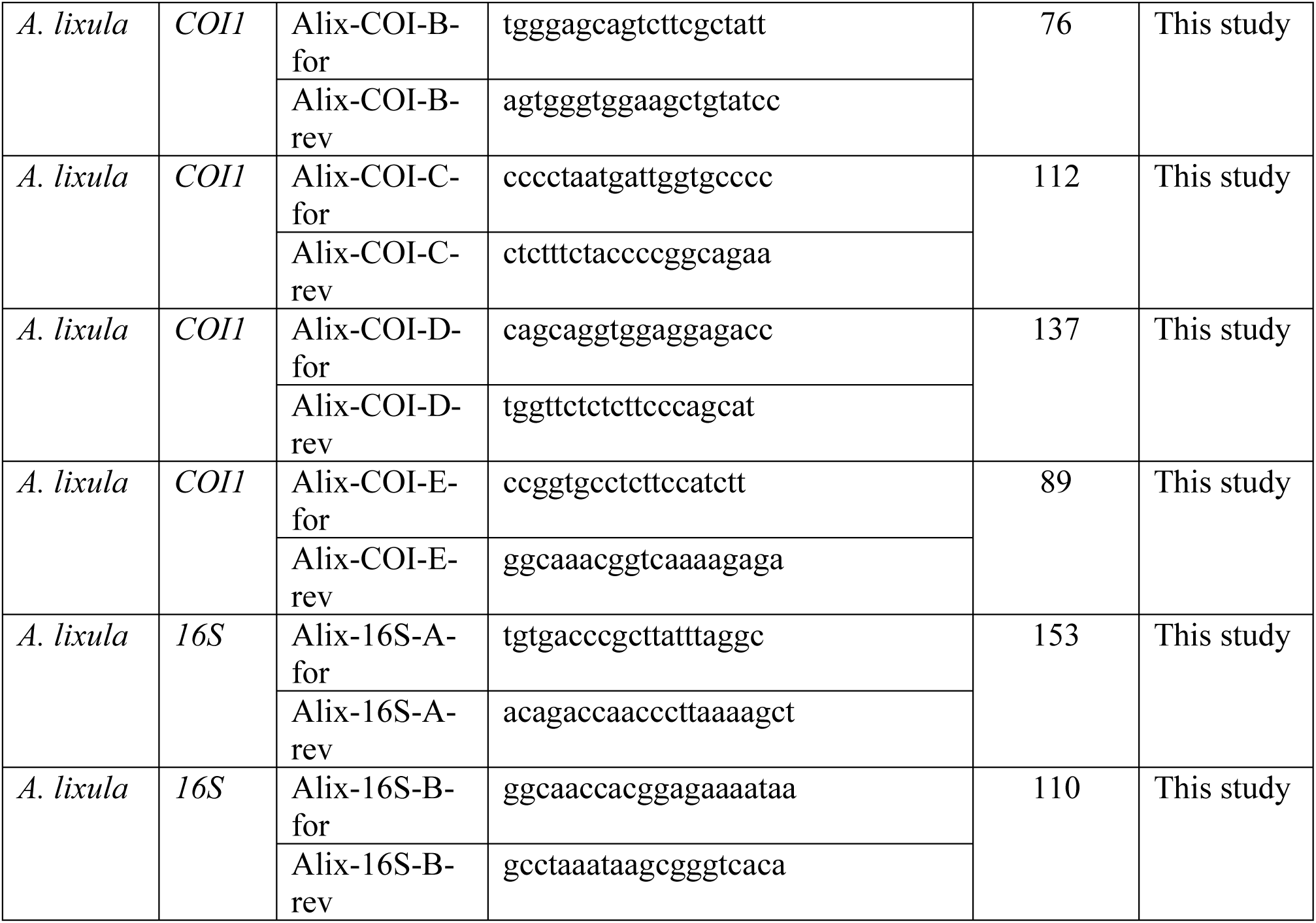
Primers used in this study.

As a test for DNA quality of the sample, we also used the primer pair jgLCO1490/ jgHCO2198 (Table 1), that was shown to robustly amplify a large range of marine invertebrate taxa(Geller et al., 2013).

### Amplification

In order to minimize the risk of cross-contamination, pre-packed, aerosol-resistant and DNA-free pipette tips were used for assembling PCR reactions and loading samples in gels. Different pipette sets and laboratory room were used when handling urchin specimens and their DNA. Pre-PCR and post-PCR setups were located in different rooms, as recommended in forensic studies (King et al., 2008).

PCR reactions were assembled with 0.4 µM each primer, 1% w/v BSA and 2 units MyTaq polymerase (Bioline) in a reaction volume of 25 µL. Amplifications were performed in 96 well plates. PCR conditions were: initial denaturation 3 min at 95°C, followed by 35 cycles of denaturation 60 sec at 95 °C, annealing 30 sec at 56 °C, extension 25 sec at 72 °C; a final extension step was set for 10 min at 72 °C. Templates in the specificity test PCR included DNA extracted from gonads of adult *P. lividus* and *A. lixula* (0.1 ng), and from invertebrates of different taxonomic groups (100 ng). In particular, we extracted DNA from the Sipunculid *Phascolosoma granulatum*; the crab *Pilumnus hirtellus*; the shrimp *Synalpheus gambarelloides*; the Ophiuridae *Amphipholis squamata*; a Terebellidae *Thelepus hamatus*; the whelk *Euthria cornea*. With the exception of the Ophiuridae and the Polychaete, either too small or unstructured to be dissected, we extracted DNA from appendices of invertebrate samples, with the aim to minimize contamination with DNA from other species contained in the gut. Templates in the sensitivity tests consisted of DNA from adults of either *P. lividus* or *A. lixula*, diluted in water from 10 ng to 10 fg (10^6^ folds dilution). For PCRs testing the inhibition effect of excess predator DNA, templates contained 0.1 ng urchin DNA and 100 ng (1,000 fold) of predator DNA. For screening all samples for presence of urchin DNA, we used 50 ng DNA as template in each reaction, as previously done in a similar survey (Redd et al., 2014). Each reaction plate contained one positive control (0.1 ng *P. lividus* DNA) and 4-5 negative controls (reaction mixture with no template).

For *COI1* amplification with Geller’s universal primers jgLCO1490/ jgHCO2198, PCR conditions were: initial denaturation 3 min at 95°C, followed by a touch down setup with 35 cycles of denaturation 60 sec at 95 °C, annealing 60 sec at 52-48 °C (-1 °C /cycle for the first four cycles, followed by 48 °C for 31 cycles), extension 60 sec at 72 °C; a final extension step was set for 10 min at 72 °C.

PCR products were loaded in 2% agarose gel and visualized by GelRed (Biotium) staining. Band presence and intensities were automatically detected by the software GelAnalyzer 23.1.1 and manually checked.

### Sequencing

Bands amplified by primers were manually excised from the gels and the DNA extracted by using the Gel Extraction Kit (Ezna, VWR) according to the manufacturer’s instructions. The purified DNAs were directly sequenced by forward primers (Table1). Sanger sequencing was performed by Eurofins Genomics (Germany). The resulting sequences were blasted in the NCBI database (BLASTn) to retrieve the identity of the specimens. Additional taxonomic information for each species (Class, Family) was obtained by the World Register of Marine Species (WORMS https://www.marinespecies.org).

### Next generation sequencing

A thorough amplicon sequencing of the locus *COI1* in invertebrate samples was performed with the Illumina-tagged primers mICOIintF and jgHCO2198, which amplify a short fragment of around 313 bp, optimal for NGS sequencing with MiSeq platform (Illumina) (Leray et al., 2013) (Table1). PCR reactions were assembled with 100 ng template, 0.3 µM each primer, 1% w/v BSA and 4 units MyTaq polymerase (Bioline) in a reaction volume of 50 µL. PCR conditions were modified from (Leray et al., 2013): initial denaturation 3 min at 95°C, followed by 16 touchdown cycles of denaturation 45 sec at 95 °C, annealing 1 min at 62-46 °C (-1 °C/cycle), extension 1 min sec at 72 °C; 25 more cycles with annealing temperature fixed at 46 °C; a final extension step was set for 10 min at 72 °C. For each sample, five replicate amplifications were pooled and purified together in order to reduce stochastic misrepresentation of rare templates. Bands of the expected size were gel-purified as described above and sequenced through MiSeq platform by Eurofins Genomics. Primer trimming and merging of paired end reads were performed by Eurofins Genomics through Cutadapt and FLASH softwares, respectively. Libraries were then processed by FROGS to remove chimera, cluster the OTUs and assign taxonomy, based on Midori’s marine database (Escudié et al., 2018).

## Results

### Invertebrate assemblages

A total of 360 individuals of potential predator invertebrates were collected and visualized at a stereomicroscope for taxonomic identification (Supplemental Table S1). Pictures of each invertebrate are shown in the Supplemental Fig S1. Only for ∼22% of specimens we were confident enough to assign an identification at the species level, especially for crustaceans. For other samples, we only reached the Genus (∼18%), Family/Superfamily (∼19%), Order/Infraorder (∼14%) or even just the Class (∼27%) taxon level. Overall, most of the collected invertebrates were crustaceans (Malacostraca, ∼40%). Brittle stars (Ophiuridea) and worms (Polychaeta) were also abundant (∼29% and ∼23% of samples, respectively). The other invertebrates belonged to Gastropoda (∼4%) and Sipuncula (∼4%) (Fig 1a).

**Fig 1.**
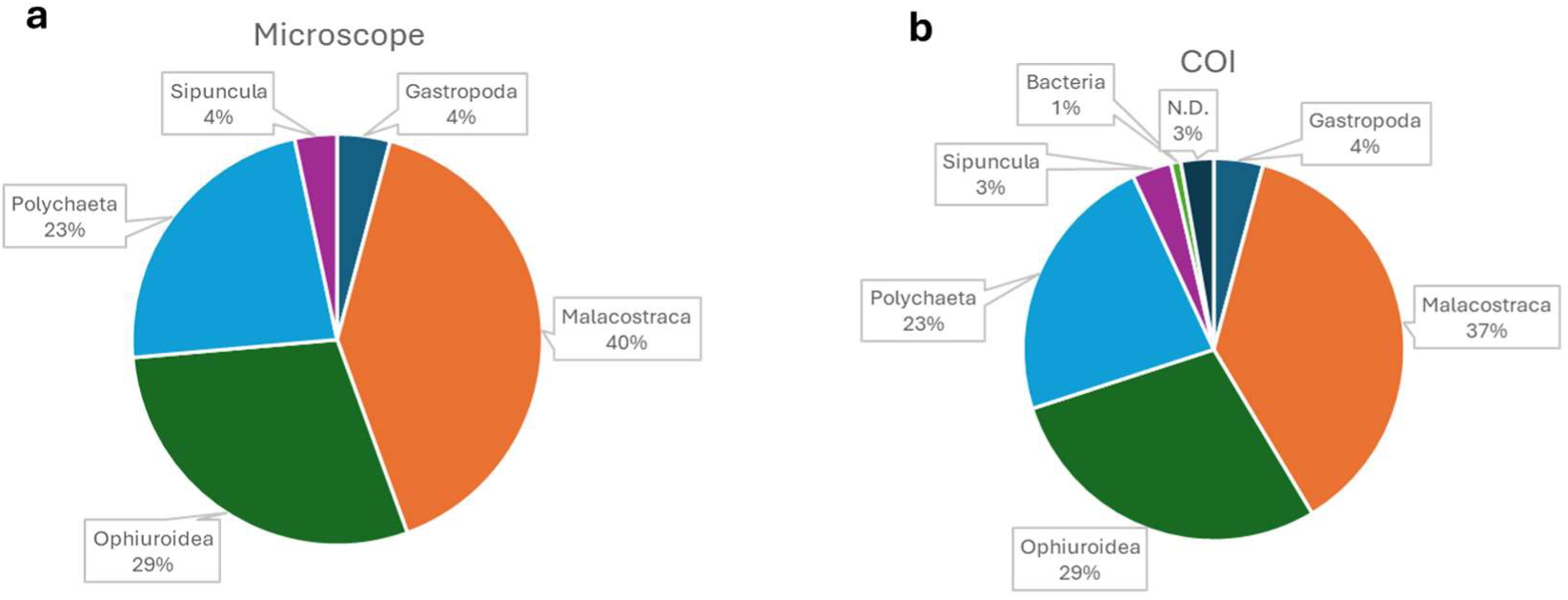
Distribution of collected samples among Classes according to visual assignment (a) and *COI1* barcode (b). N.D. = not determined.

### DNA quality and barcoding

To check for integrity of the DNA extracted from preserved invertebrate samples, we analyzed a subset of randomly selected samples by electrophoresis. Most samples showed an extended smear of low molecular weight DNA, indicative of degradation (Supplemental Fig S2). Very little high molecular weight genomic DNA was visible. Degradation was likely a result of sample storage and/or handling during the extraction protocol. Concerned about the quality of the samples, we first tested all 360 potential predators for their ability to support PCR amplification. To this end, we used Geller’s “universal” *COI1* primer pair that had been shown to amplify a wide range of invertebrate taxa (Geller et al., 2013). Accordingly, the vast majority (∼97%) of samples gave a clear and strong band, indicating that the handling and storage of the animal and the extraction protocol did not affect the ability of the DNA to amplify (Supplemental Figure S3). Only 10 invertebrate samples failed to amplify (Supplemental Table S1). We excluded that the PCR failure was due to incompatibility of the primer set with the sample species, since other members of the same species showed a clear band in other reactions. More likely, the negative result for these 10 samples was due to the presence of PCR inhibitors, or to insufficient DNA quality. In all the other 350 invertebrate samples clear bands were visible with size ranging between 400 and 1,000 bp, indicating that the recovered DNA was not fully degraded, and that sufficient amount of nucleic acid was long enough to serve as template (Supplemental Figure S3). Since the primers for urchins were designed on a much smaller region (< 200 bp) and for genes encoded in the mtDNA, present in high copy number, we were confident that our strategy had the potential to work even when the target DNA was present in trace amounts.

The bands amplified in the *COI1* region by Geller’s primers were purified and sequenced to serve as barcode for molecular identification of invertebrates and urchins. Out of 350 invertebrate sequences, three failed to provide a reliable identification, due to misassignment to Bacteria. Supplemental Table S1 reports the names of species most closely related to sequences for each sample and the identity scores. Most of sequences (63%) retrieved a high confidence hit (identity ≥ 95%). Samples with the lowest identity confidence (75-85%) were especially abundant among Polychaetes. Overall, molecular barcoding confirmed the coarse visual assignment and provided a finer labeling. At the Class level, the molecular identification agreed with visual assignment for all samples. At the Family level, 14 samples (4%) showed a disagreement between *COI1* sequencing and visual assignment.

Class distribution was similar to the one obtained by visual assignment (Fig 1b); minor deviations were due to the 13 specimens that failed to provide a valuable barcode sequence.

### Barcoding of urchin juveniles

In the study area, the two most abundant sea urchin species are *P. lividus* and *A. lixula*. Adults are easily recognized by color, form and behavior. However, small juveniles (< 5 mm) are indistinguishable, since spines are still colorless and the shape of the urchin is uniform. In our surveys we collected 55 sea urchin juveniles. Most of them (45, ∼82%) were smaller than 1 mm, the rest being sized between 1 and 5 mm (Supplemental Table S2). In order to identify the urchin species, we sequenced the *COI1* loci of a subset of 37 urchins (Supplemental Figure S2). Only two samples failed to provide a reliable identification, with one sequence with no significant similarity in databases, and the other sequence assigned to Bacteria (Supplemental Table S2). The remaining 35 juveniles were all assigned to *P. lividus*, suggesting that the settlement event that spanned our surveys was attributed to that species.

### Primer efficiency

Ideal primer sets for assessing predator-prey interaction strongly amplify the prey DNA but not other organisms’ DNA. In order to design primers specific for *P. lividus*, yet able to amplify *P. lividus* accessions from different Mediterranean regions, we aligned 254 mtDNA sequences of *P. lividus* from different Mediterranean areas. To ensure specificity, the alignment included 328 sequences of the sympatric sea urchin species *A. lixula* and those of their potential predators, belonging to different taxonomic groups. We then selected the genetic regions that were conserved within the same species, but that differed between the two sea urchin species, and also with other invertebrates. In particular, for *P. lividus* we designed five primer pairs for the locus of the *cytochrome oxidase c subunit 1* (*COI1*), three pairs for *cytochrome oxidase b* (*CYTb*) and one pair for the *16S* locus; for *A. lixula*, we designed five pairs for *COI1* and two pairs for *16S* (Table 1). Each primer pair was tested for specificity of amplification in a panel comprising positive controls, i.e. genomic DNA of adult *P. lividus* and *A. lixula*, and negative controls, chosen among potential predator taxa. The expected band sizes were in the 63-178 bp range, in order to amplify degraded DNA.

We first tested for primer specificity- Primer sets amplified urchin DNA with different efficiencies, as observed by band intensity (Supplemental Fig S4). For *P. lividus* specific primers, all primers produced a strong band with *P. lividus* DNA, with the exception of Pliv-CYTb-B, which failed to amplify. Pliv-COI-D and Pliv-COI-E showed unspecific bands in the predator samples. Conversely, Pliv-COI-A, Pliv-COI-B, Pliv-COI-C, Pliv-CYTb-A, Pliv-CYTb-C and Pliv-16S, only showed the specific band when *P. lividus* DNA was present in the template. The short 63 bp amplicon of Pliv- COI-A, however, was faint and too close to the primer bands. For this reason, we excluded Pliv-COI- A from further analyses. For *A. lixula*, all primers produced a band in presence of *A. lixula* DNA. However, Alix-COI-A, Alix-COI-D, Alix-COI-E and Alix-16S-B also produced unspecific bands in presence of predator DNA. Only Alix-COI-B, Alix-COI-C and Alix-16S-A were specific. For their specificity in the reaction, we used primers Pliv-COI-B, Pliv-COI-C, Pliv-CYTb-A, Pliv-CYTb-C and Pliv-16S, as *P. lividus* specific primers, and Alix-COI-B, Alix-COI-C and Alix-16S-A as *A. lixula* specific primers in all subsequent experiments.

Once we selected specific primers for *P. lividus* and *A. lixula*, we tested for their sensitivity, by diluting urchin DNA from 1 ng to 10 fg for *P. lividus* and from 10 ng to 100 fg for *A. lixula* (100,000 fold). A strong band was visible for all primer pairs with as little as 10 pg DNA, and fainter bands, yet detected by the automatic image analyser, up to 1 pg for Pliv-COI-B and Pliv-16S (Supplemental Fig. S5; Supplemental Table S3). The selected primers therefore are sensitive enough to detect the small amount of urchin DNA present in the stomach of predators. Our tests were performed on purified DNA. However, in homogenates from whole predators and their gut content, the presence of excess amount of predator’s DNA might interfere with prey’s DNA amplification. To test this hypothesis, we diluted a small amount (100 pg) of urchin DNA with a thousand-fold excess of invertebrate DNA, selected from a representative of each major taxonomic group. All the primers amplified the corresponding urchin DNA with no interference from invertebrate DNA (Supplemental Fig S6), indicating that they are suitable for detecting the prey within the predators’ body.

### Detection of urchin DNA in invertebrate samples

The 55 sea urchin juveniles all strongly amplified with *P. lividus* primers Pliv-COI-B, Pliv-CYTb-A and Pliv-16S, as evidenced by the intense bands of the expected sizes (Supplemental Fig S7). Conversely, the same samples did not produce any band when *A. lixula* specific primers Alix-16S-A were used. This result confirmed the barcoding results, indicating that all the juveniles belonged to the species *P. lividus*. Next, we then tested the invertebrates DNA with the the five *P. lividus* specific primers Pliv-COI-B, Pliv-COI-C, Pliv-CYTb-A, Pliv-CYTb-C and Pliv-16S. The 10 invertebrate samples that had failed in the amplification with Geller’s universal primers also did not amplify with *P. lividus* primers, confirming that they might contain PCR inhibitors or that their DNA quality was too low. Among the remaining 350 samples, 30 (9%) scored positive for all five loci, 54 samples (15%) for at least four loci, 73 (21%) for at least three loci. Additional 47 (13%) and 84 (24%) samples showed a band for just two or one locus, respectively. Conversely, 145 samples (41%) did not show any band for any primer set (Fig 2a and Supplemental Table S1). Pliv-16S was the most sensitive locus, with 149 positive samples (43%), followed by Pliv-CYTb-A (n=121, 35%), Pliv-COI-C (n=95, 27%), Pliv-CYTb-C (n=64, 18%) and Pliv-COI-B (n=52, 15%). Excised and gel-purified bands from eight positive samples and the positive control (adult *P. lividus*) amplified with the Pliv-16S, Pliv- CYTb-A and Pliv-COI-B primer pairs were sequenced. Though short, all sequences belonged to *P. lividus*, confirming that the DNA of this urchin species was present in the samples (Supplemental Table S4).

**Fig. 2.**
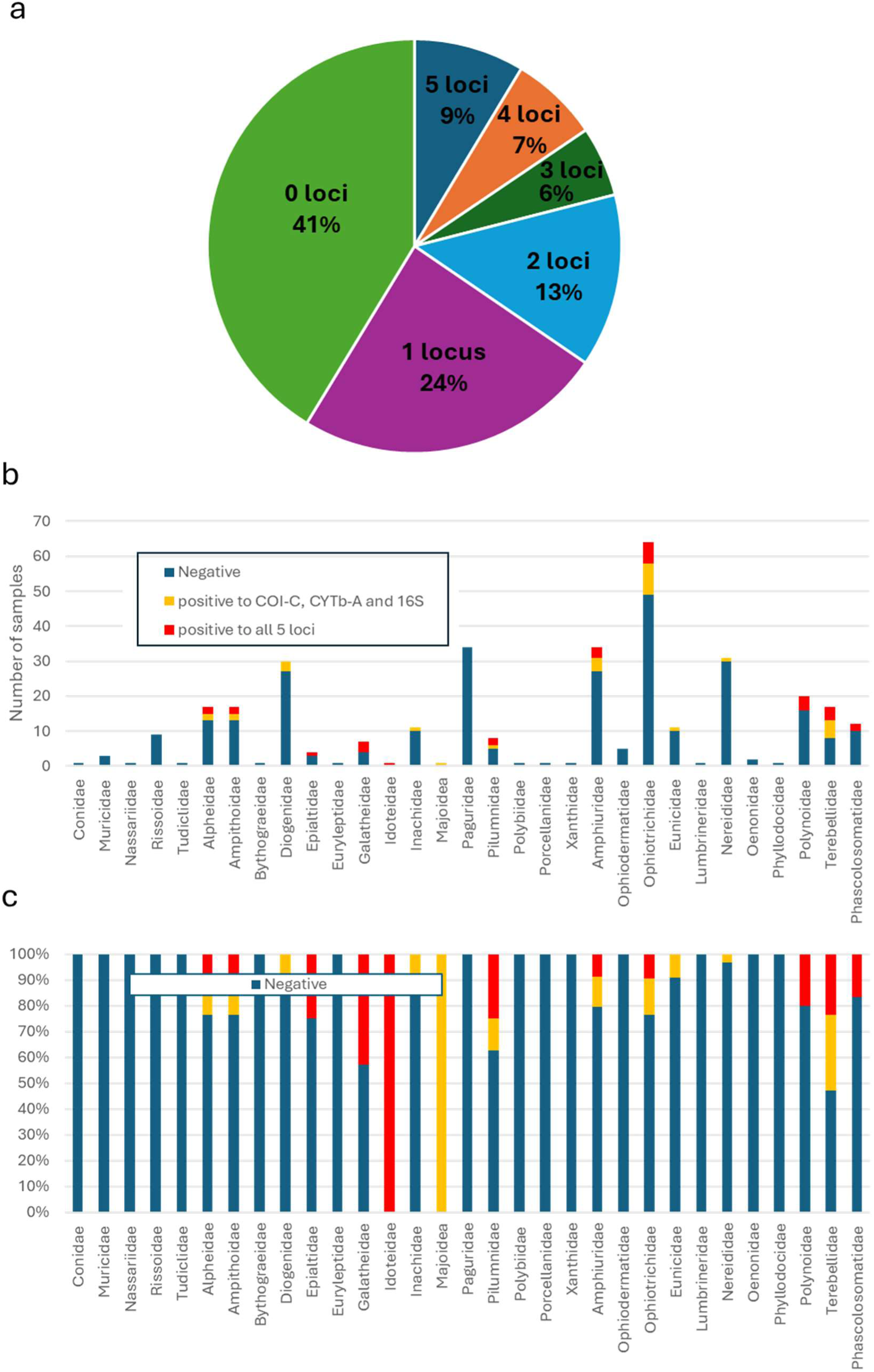
Distribution of samples showing positive response to 0-5 P. lividus loci (a). Proportion of samples positive to all five loci or for the three best performing primer pairs, over the total collected samples, expressed as number of samples (b) and percentage (c) within each Family.

Considering only the most conservative results from the 30 samples that strongly amplified all five *P. lividus* loci, positive invertebrates belonged to all Classes but Gastropoda, with 11 positive Malacostraca (8% of all samples from the same Class), nine Ophiuroidea (9%), eight Polichaeta (10%) and two Sipuncula (17%) (Table 2). Among Families, the rate of positive invertebrates ranged widely. Particularly represented were shrimps from Alpheidae and Ampithoidae (both n=2, 12%); crabs from Galatheidae and Pilumnidae (n=3 and 2, 43% and 25%, respectively); brittle stars from Ophiotrichidae and Amphiuridae (n=6 and 3 respectively, 9% each); worms from Polynoidae and Terebellidae (n=4 each, 20% and 24% respectively) (Table 2; Fig 2b, c). If we do not consider the two loci which underperformed, Pliv-CYTb-C and Pliv-COI-B, the number of samples positive in all three remaining loci was 60 (17%), distributed among Malacostraca (n=21, 16%), Ophiuroidea (n=22, 21%), Polichaeta (n=15, 18%) and Sipuncula (n=2, 17%) (Table 2). The represented Families were similar to the distribution with five loci, but with more samples, especially among the shrimps Alpheidae and Ampithoidae, which doubled their representation, the brittle stars Ophiotrichidae and Amphiuridae and the worms Terebellidae, which increased even more (n=15, 7 and 9, respectively, corresponding to 23%, 21% and 53%). Few positive samples were also present among the crabs Diogenidae, Inachidae and Majoidea, and the worms Eunicidae and Nereididae (Table 3; Fig 2b, c).

**Table 2.**
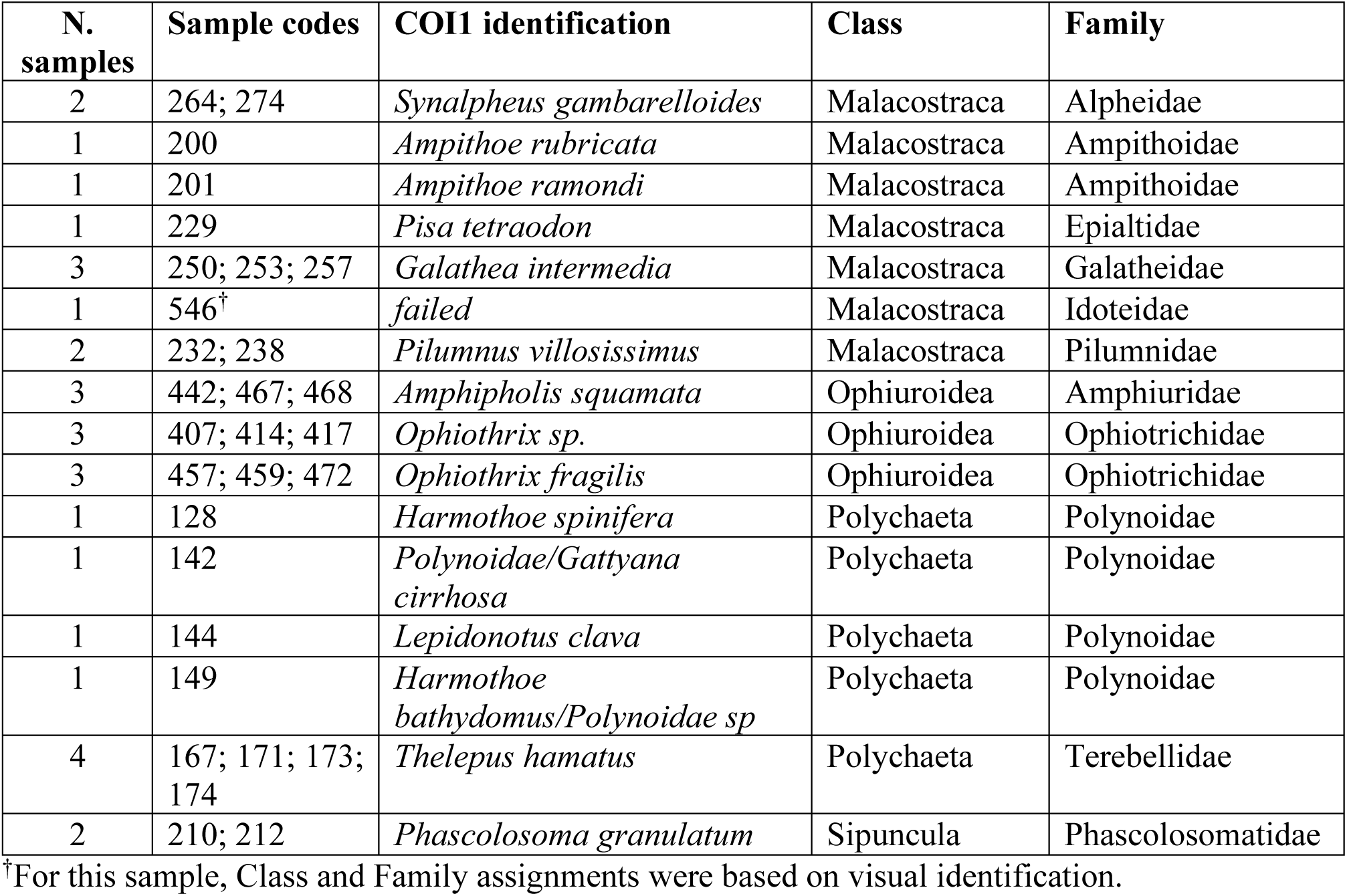
List of samples positive to all five *P. lividus* loci.

**Table 3.** Additional samples that scored positive to the three best performing primer pairs Pliv-COI- C, Pliv-CYTb-A and Pliv-16S.

| N. samples | Sample codes | COI1 identification | Class | Family |
| --- | --- | --- | --- | --- |
| 2 | 265; 268 | *Synalpheus gambarelloides* | Malacostraca | Alpheidae |
| 2 | 195; 197 | *Ampithoe rubricata* | Malacostraca | Ampithoidae |
| 3 | 345; 354; 545 | *Calcinus tubularis* | Malacostraca | Diogenidae |
| 1 | 245 | *Inachus aguiarii* | Malacostraca | Inachidae |
| 1 | 248 | *Herbstia condyliata* | Malacostraca | Majoidea |
| 1 | 235 | *Pilumnus hirtellus* | Malacostraca | Pilumnidae |
| 4 | 443; 447; 448; 470 | *Amphipholis squamata* | Ophiuroidea | Amphiuridae |
| 6 | 406; 458; 461; 473; 483; 485 | *Ophiothrix fragilis* | Ophiuroidea | Ophiotrichidae |
| 3 | 408; 409; 415 | *Ophiothrix sp.* | Ophiuroidea | Ophiotrichidae |
| 1 | 141 | *Eunicidae sp/Palola* | Polychaeta | Eunicidae |
| 1 | 127 | *Platynereis sp* | Polychaeta | Nereididae |
| 5 | 165; 166; 168; 169; 170 | *Thelepus hamatus* | Polychaeta | Terebellidae |

### Next Generation Sequencing of COI1 assemblages

As a further, independent test for the presence of urchin DNA within the invertebrate bodies, we performed a full Next Generation Sequencing (NGS) of the *COI1* locus amplified with Leray’s universal primers for a subset of invertebrate samples, namely three brittle stars (*Ophiothrix* sp.), three shrimps (*Synalpheus gambarelloides*), and two crabs (*Galathea intermedia*) (Table 4). As expected, the majority of reads (53-96%) within each sample belonged to the host organism. Yet, sequencing of *COI1* amplicons for all the specimens which had showed a strong band with all five *P. lividus* primers, also revealed the presence of *P. lividus* reads in their full COI1 assemblages (0.02- 0.5%), with high identity score (>94%). Conversely, the brittle star, shrimp and crab samples which had resulted negative with *P. lividus* primers, did not present any urchin COI1 read.

**Table 4.** NGS *COI1* results for a subset of samples.

| Sample code | Organism | Positivity to *P. lividus* primers | Host reads (%) | *P. lividus* reads (%) | Minimum *P. lividus* identity (%) |
| --- | --- | --- | --- | --- | --- |
| 408 | Ophiotrix sp. | yes | 96,03 | 0,24 | 98,81 |
| 409 | Ophiotrix sp. | yes | 70,81 | 0,02 | 99,68 |
| 410 | Ophiotrix sp. | no | 95,59 | 0 |  |
| 274 | *S. gambarelloides* | yes | 80,85 | 0,5 | 99,68 |
| 264 | *S. gambarelloides* | yes | 96,07 | 0,07 | 97,75 |
| 269 | *S. gambarelloides* | no | 52,66 | 0 |  |
| 253 | *G. intermedia* | yes | 92,59 | 0,02 | 94,6 |
| 252 | *G. intermedia* | no | 95,6 | 0 |  |

## Discussion

Sea urchins play a pivotal role in driving infralittoral system dynamics, with high-density populations exerting a bulldozing effect on macroalgae, leading to barren formation (Steneck, 2013). Estimates of predation pressure on sea urchins suggest a bimodal distribution related to urchin size, with small juveniles hiding efficiently in crevices, and adult urchins may avoid predation due to their size and defense spines. According to this model, intermediate sized urchins are more susceptible to predation (Fagerli et al., 2014; Ling & Johnson, 2012; Sala & Zabala, 1996; Shears & Babcock, 2002; Tegner M.J. & Dayton P.K., 1981). However, these estimates primarily consider predation by fishes and large invertebrates such as lobsters. Smaller, crawling invertebrates might efficiently extract urchin juveniles from crevices, exerting the most prominent control of urchin populations (Bonaviri et al., 2012; Clemente et al., 2013; McNaught Douglas Colin, 1999; Pavlova, 2009; Scheibling & Hamm, 1991; Scheibling & Robinson, 2008; Smith et al., 2023; Steneck et al., 2013). Accordingly, for the Mediterranean urchin species *P. lividus*, over 75% of juveniles disappear within six months from settling in algalforests (Sala & Zabala, 1996), whereas they persist in greater numbers in barrens. Compared to barren, forests host a large abundance and variety of invertebrates, with up to 200 species per m^2^ (Christie et al., 2003). In our surveys, we collected 360 invertebrates from Mediterranean forests, belonging to Malacostraca, Ophiuroidea, Polychaeta, Gastropoda and Sipuncula. Species assignment requires a thorough level of taxonomic expertise, with scholars usually specializing in one or a few Classes, and/or limited geographical regions. Moreover, taxonomic knowledge is a field suffering a severe decline in practitioners (Drew, 2011). Visual identification of a wide collection of invertebrates is, therefore, a daunting task requiring the involvement of multiple, uncommon experts. When samples are preserved for a long time or are not carefully handled, they may also lose structural details (such as color and appendages) that make taxonomic identification more challenging. Accordingly, several of our samples presented fragmented bodies (Supplemental material S1), due to processing of samples and their transport from the field to the dissecting and molecular laboratories, located in different countries. By microscopic analysis, we were able to identify only 78 specimen (22%) at the species level, for the vast majority of which were crustaceans (46% of all samples from Class Malacostraca). Our taxonomic confidence was lower for Ophiuroidea (8%), whereas we did not assign a specific name to any of molluscs, Sipuncula or Polychaeta. For over 27% of samples, we even limited our identification to the Class level, especially for Polychaeta (77%) and Ophiuroidea (29%). Due to our lack of taxonomic expertise for many organisms, we relied on molecular barcoding as an aid to species identification (Antil et al., 2023). Over the years*, cytochrome c oxidase 1* (*COI1*) has become the standard locus for barcoding of Metazoans, with thousands of annotated sequences deposited in public databases, such as BOLD (Barcode of Life data system). In order to sequence the *COI1* locus, we first had to verify that the stored and processed samples were able to deliver DNA of sufficient quantity and quality for amplification. Despite most samples failing to show a high molecular weight band upon electrophoresis, indicative of abundant and intact genomic DNA, and many samples even presented a smear of degraded DNA, *COI1* amplification and sequencing were successful for over 97% of samples (Supplemental Table S1). Only 10 samples failed to produce a visible band (Supplemental Fig S3), probably because their DNA was too degraded to amplify, or because the extract contained PCR inhibitors. Consequently, although the same samples were also negative when tested with *P. lividus* specific primers, we cannot draw any conclusion regarding the presence of urchin DNA. For additional three samples, *COI1* sequencing resulted in a *Photobacterium* spp. assignment rather than the related crabs. *Photobacterium* species are marine microorganisms, some known as pathogens for fishes and crabs (Rivas et al., 2013), but they can also constitute the major component of crab microbiome (Jiang et al., 2023). It is therefore possible that these three specimens were infected by the bacterium, whose DNA abundance predominantly outcompeted the amplification reactions. Overall, barcoding strongly correlated with visual assignment. Minor disagreement at the Family level (4%) was most likely due to visual misidentification. Though generally considered a potent aid in taxonomic identification, it must be noted that molecular barcoding may suffer from inherent limitations. First, it relies on sequence comparison of a single locus of mtDNA, characterized by a high mutation rate and no recombination. As a result, geographically or sexually separated populations of the same species may diverge in sequence (intraspecific polymorphism), leading to an overestimation of the number of effective species(Eberle et al., 2020). Conversely, for some taxa *COI1* sequence is not variable enough to discriminate between morphologically different species (Carew & Hoffmann, 2015). Finally, a bias in the sequence deposition into repositories leads to overrepresentation of species from certain areas of the world. In our survey, most of the invertebrates (63%) were assigned to species with high confidence (>95% identity). For the specimen with lower identity score, it is safer to restrict the identification to higher taxonomic levels, such as Genus or even Family. For example, all our Terebellidae samples were assigned to *Thelepus hamatus* with an identity score of 88-89%. *T. hamatus* is distributed along the Pacific coast, and clearly it is not the correct assignment for our sample. It is therefore likely that the *COI1* sequence of the related Mediterranean species is not represented yet in the reference databases, and its correct assignment was missed.

Full *COI1* Next Generation Sequencing may also be used to infer the presence of all species present in a sample, with their relative abundances (Clare, 2014; Pompanon et al., 2012). However, each analysis is costly and time consuming. For a large set of samples, or when only one or few preys need to be identified, the direct PCR assay with specific primers is a more affordable and rapid method of detection. In our study, we used NGS amplicon *COI1* sequencing for a subset of eight samples, to confirm the validity of our specific primer sets in detecting urchin DNA.

In the Mediterranean Sea, shallow rocky reefs are dominated by two sympatric species, *P. lividus* and *A. lixula*. At the settler stage, these urchins are indistinguishable. Sequencing of *COI1* locus for a subset of the collected settlers revealed that they all belonged to *P. lividus*. Yet, we designed and tested specific primers for both sea urchin species, in order to provide tools for molecular trophic studies in future spawning events in the Mediterranean and Atlantic. For primer design, we focused on mitochondrial DNA (mtDNA), since each diploid cell contains only two copies of nuclear loci, but thousands of circular mtDNA molecules (Zhang Yaping et al., 1993), which are less prone to degradation, making it easier to detect in digested samples. Additionally, sequences of mtDNA loci are widely available in public database for a large variety of species, facilitating the design of prey- specific primers. We selected three loci within the mtDNA: *COI1*; *CYTb*; and *16S*. *COI1*, being the barcode standard, is highly polymorphic and sequences are available for a large number of individuals within species, accounting for intraspecific variability. *COI1* is also the most commonly used locus for assessing trophic relationships among species, reviewed in (King et al., 2008; Pompanon et al., 2012). *CYTb* has been used for Insects, Arachnida and fishes (Parsons et al., 2005; Pons, 2006). Additionally, *CYTb* sequences have been used to discern the phylogeographic distribution of *P. lividus* populations in the Mediterranean (Maltagliati et al., 2010). *16S* locus is also polymorphic, and among other species, it has been used to detect DNA of the sea urchins *Centrostephanus rodgersii* and *Heliocidaris erythrogramma* is lobster fecal samples in Tasmania (Redd et al., 2014; Smith et al., 2023).

Due to PCR sensitivity, caution is required when performing and interpreting molecular analyses, with multiple tests and controls needed (reviewed by (King et al., 2008)). The specific primers for *P. lividus* and *A. lixula* were designed by aligning hundreds of accessions of both urchin species from across the Mediterranean and Atlantic. Since these primers amplified DNA from urchins collected both in Catalonia and Sicily, located in distant areas of the Mediterranean, they proved not to be affected by intraspecific genetic variability. Primers were also tested for specificity, both in silico and by PCR, including invertebrates encompassing the taxa collected in our surveys. We then selected five primer pairs specific for *P. lividus* and three primer pairs specific for *A. lixula*. Sensitivity tests showed that these primers were able to detect as little as 1-10 pg urchin total DNA. With a nuclear genome of 927.4 Mb and 2,000-20,000 copies of 15.697 Kb mtDNA per cell, 1 pg total DNA corresponds to about 1 single cell, or 1,400-20,000 mtDNA copies (Cantatore et al., 1989; Marlé Taz et al., 2023). However, estimates in samples with degraded DNA may be skewed. Primer efficiency was also not affected by a thousand fold excess of exogenous DNA, from invertebrates belonging to different Classes, mimicking the scenario occurring in micropredators feeding on urchin settlers. Having passed all the control tests, the selected primer pairs can be confidently used as a molecular tool to study trophic interactions involving the sea urchins *P. lividus* and *A. lixula*.

Quantitative assessment of dietary intake in predator-prey interactions is a challenging task. Molecular techniques have been used to quantify the amount of prey consumed in captivity trials (Deagle & Tollit, 2007), but evaluation from field samples is not straightforward (Nejstgaard et al., 2008; Redd et al., 2014; Troedsson et al., 2007; Weber & Lundgren, 2009). The amount of prey DNA is dependent on the time of ingestion and, in the case of different predator species, on their rate of digestion. Therefore, it is usually not possible to discern between low levels of recent ingestion and high levels of past ingestion, when most of the DNA has been degraded. For this reason, it is safer to adopt a qualitative approach with a binary outcome, positive or negative, regarding the presence of prey DNA. It must be stressed, however, that also qualitative assessments are subject to a certain degree of subjectivity in the choice of the threshold separating positive from negative samples. Threshold values can be based on the band intensity, as in this and other studies (Smith et al., 2023), or on Ct values in the case of qPCR analyses (Redd et al., 2014). In the present work, we relied on the automatic detection of bands, over the gel background, by specific software. To be more conservative in the result interpretation, the threshold was set high enough to discard the ambiguous bands, visible but of faint intensity. Though this is a common approach, it still retains an arbitrary component that needs to be taken into account. Additionally, the choice of small amplicon sizes, instrumental in detecting degraded prey’s DNA in gut content, presents the risk of spurious bands resulting from primer dimerization, which may affect interpretation. Longer electrophoresis separation and careful size interpolation may facilitate the discrimination between genuine and artifactual bands. In our analyses, the largest share of samples (41%) was negative, showing no visible bands. Therefore, the issue of dimer bands probably does not apply to our amplification conditions, as it would affect samples regardless of the presence of template. As an additional measure of confidence in result interpretation, we decided to analyze multiple loci. For *COI1* and *CYTb*, we also used two primer pairs spanning different regions of the loci. The number of positive bands was correlated with the amplicon size. Pliv-COI1-C, Pliv-CYTb-A and Pliv-16S showed the highest number of positive samples, and their amplicon sizes were 178, 155 and 161 bp, respectively. The amplicons Pliv-COI-B and Pliv-CYTb-C, in contrast, were present in fewer samples, and their sizes were smaller (93 and 119 bp, respectively). It is possible then that the higher stringency of Pliv-COI- B and Pliv-CYTb-C derives from the more difficult discrimination of amplicon bands from primers. The assignment of the amplified bands of all three loci to *P. lividus* DNA was further confirmed by sequencing. It is very important to stress that direct predation is not the only event leading to the presence of urchin DNA in samples. Contamination is an obvious reason, though a careful handling procedure, use of separate machinery for samples and standard *P. lividus* DNA, and inclusion of several negative controls in each PCR run, let us to believe that we prevented contamination to occur in our analyses. Accordingly, the largest share of samples (41%) did not amplify at any locus. Urchin DNA can be present within an invertebrate body as a result of secondary predation, i.e., when a predator eats another predator that had fed on *P. lividus*. Another level of trophic interaction that cannot be excluded by molecular analyses is scavenging on dead urchin material, rather than direct predation on settlers. Finally, urchin DNA can be present in the environment through their feces or disintegration of their remains. This aspect is particularly concerning, since direct testing of unconsolidated sediments in Tasmanian reefs found urchin DNA in 20-100% of samples (Redd et al., 2014), and sediments are known to be repositories for eDNA (Bowman & McCuaig, 2003). However, only a minority of lobsters fed with those sediments or directly with urchin feces scored positive, probably because digestion eliminated the already degraded urchin DNA present in the sediment and urchin feces, suggesting that environmental DNA might represent a minor issue (Redd et al., 2014). Accordingly, a large portion of our samples was negative. Yet, it is possible that the presence of P. *lividus* DNA in some of our positive samples derives from sediment or particulate feeding. This is especially likely for the Terebellidae and Sipuncula specimens, known to feed by sediment filtration and accumulate organic material. All the other positive invertebrates are carnivorous, and events of direct predation on *P. lividus* settlers are the most likely scenario. Members from the Alpheidae, Pilumnidae, and Galatheide families among crustaceans were particularly abundant as positive samples, especially relative to the total number of specimens of the same taxa collected (24%, 38% and 43%, respectively). Earlier works in aquaria showed that *Alpheus dentipes*, *Pilumnus villosissimus* and *Pilumnus hirtellus* are able to feed on *P. lividus* settlers (Bonaviri et al., 2012). Moreover, field surveys in the Atlantic observed a negative correlation between the abundance of sea urchin settlers and that of *Alpheus macrocheles*, and other decapods (Cano et al., 2024; Williams et al., 2011). In our study, we found four Alpheidae members (24% within the Family) and three *P. villosissimus* (38%) positive for *P. lividus* DNA. Crabs from the Xanthidae Family were major predators from aquaria tests (Bonaviri et al., 2012), but unfortunately, we only found one specimen in our survey, which resulted negative. Conflicting evidence in aquaria tests indicates hermit crabs (Paguridae) as potential predator of sea urchin juveniles, with some studies showing predation and others not (Bonaviri et al., 2012; Clemente et al., 2013; Fagerli et al., 2014; Scheibling & Robinson, 2008). In our survey, despite the Paguridae Family being largely represented, with 34 samples, none was positive. Most likely then, hermit crabs are not effective predators of urchin settlers and the predation events measured in aquaria represent artificial effects due to limited food choice and captivity conditions. In agreement with this hypothesis, *Pagurus bernardus* mostly fed on settlers of the urchin *Strongylocentrus droebachiensis* in aquaria when other food sources, such as clams, were not present (Fagerli et al., 2014). Moreover, in situ video recording showed that hermit crabs, though abundant, did not feed on tethered urchins along the Norwegian coast (Fagerli et al., 2014). It remains to be tested whether crabs from the Galatheidae Family, among the most common positive samples in our survey (43%), are effective predators in aquaria trials. Whelks were also negative, when five or the three best performing loci were considered. This finding is in agreement with other studies, that showed that the whelk *Buccinum undatum* interacted, without feeding, with tethered urchins (Fagerli et al., 2014) nor did it feed on urchin juveniles in captivity trials (Scheibling & Hamm, 1991; Scheibling & Robinson, 2008). Several positive invertebrates were found among two Families of Ophiuridae collected in our survey: Amphiuridae and Ophiotricidae (24% and 28%, respectively). Surprisingly, brittle stars have been overlooked so far as potential predators of young sea urchins, although our findings suggest they might represent a major component of this community. Among Polychaetes, we found many Polynoidae (20%) and some Eunicidae (9%) and Nereididae (3%) positive for *P. lividus* DNA. In aquaria tests, *Nereis sp.* did not feed on *S. droebachiensis* settlers (Fagerli et al., 2014; Scheibling & Robinson, 2008), although the tested size of urchin was larger than the size of *P. lividus* settlers we commonly found in our surveys, which might act as a size exclusion threshold for this Class of predators.

In conclusion, this study provides the molecular tools useful to study the trophic interactions involving the two major sea urchin species in the Mediterranean, *P. lividus* and *A. lixula*, whose population outbreak is responsible for the onset and maintenance of the barren status in rocky reefs. Our survey, conducted in two distant forest areas of the Mediterranean, confirmed the ability of some crustacean species to feed on *P. lividus* settlers, and revealed many more potential micropredator candidates among different Classes and Families, which altogether may act as the major controlling factor in sea urchin population density. Predation trials targeting these invertebrate species may confirm their potential micropredatory behavior.

## Acknowledgements

This work was supported by two grants: 1) “Agreement CNR/MoS (Montenegro) 2015-16: Identification of micropredators of barren-forming sea urchins by molecular markers” and 2) “Luigi e Francesca Brusarosco 2016 - Società Italiana di Ecologia: Molecular markers for the identification of sea urchin micropredators: barren/forest changes in Mediterranean rocky reefs”. The Galaxy server used for some calculations is maintained by the Institut Français de Bioinformatique (IFB), funded by the Programme d’Investissements d’Avenir (PIA), grant Agence Nationale de la Recherche, number ANR-11-INBS-0013.

We thank Dr. Massimo Turina (CNR, Italy) for the fruitful discussion that led to the idea of this project.

## Data Accessibility and Benefit-Sharing

The data that supports the main findings of this study are available in the supplementary material of this article. Other data are available from the corresponding author upon reasonable request.

## Author contributions

RDM and CB conceived the idea and analyzed the data. RDM designed and coordinated the research, developed the primers and wrote the manuscript. FDT and BH collected the samples. FDT, VM and OM performed the morphological identification. AS and FLB performed the DNA extraction and amplification. RDM, FDT and VM acquired the funding. All authors read and approved the final version of the manuscript.

---

Fig. S1. Pictures of collected invertebrates.

Fig. S2. Gel showing degraded total DNA extracted from randomly selected invertebrate samples.

Fig. S3. Gels showing COI1 amplification in the collected samples by universal jgLCO1490/jgHCO2198 primers. NC= negative controls. Marker: 50 bp.

Fig. S4. Specificity test. Gels showing amplification of representative samples from different taxa, including positive controls *P. lividus* and *A. lixula* and negative controls (NC). Marker: 50 bp.

Fig. S5. Sensitivity test. Gels showing amplification of *P. lividus* and *A. lixula* DNA in a dilution concentration series. NC= negative controls. Marker: 50 bp.

Fig. S6. Inhibition test. Gels showing amplification of *P. lividus* and *A. lixula* DNA when mixed with 1,000 thousand-fold excess of predator’s DNA. NC= negative controls. Marker: 50 bp.

Fig. S7. Gels showing amplification from urchin settlers’ DNA with *P. lividus* and *A. lixula* specific primers. NC= negative controls. Marker: 50 bp.

Table S1. List of collected invertebrates. The last three columns indicate the presence of amplified band with the corresponding *P. lividus* primer pair.

Table S2. List of collected sea urchin juveniles.

Table S3. Detection of bands in the sensitivity test for each primer pair for *P. lividus* and *A. lixula*. Table S4. Sequence of amplified bands from representative invertebrates with the specific *P. lividus* primer pairs.

## Notes

The authors declare no conflict of interest.

### Competing Interest Statement

The authors have declared no competing interest.

