## Supplementary material for "Unravelling hidden trophic interactions among sea urchin juveniles and macroinvertebrates by DNA amplification": S2-7

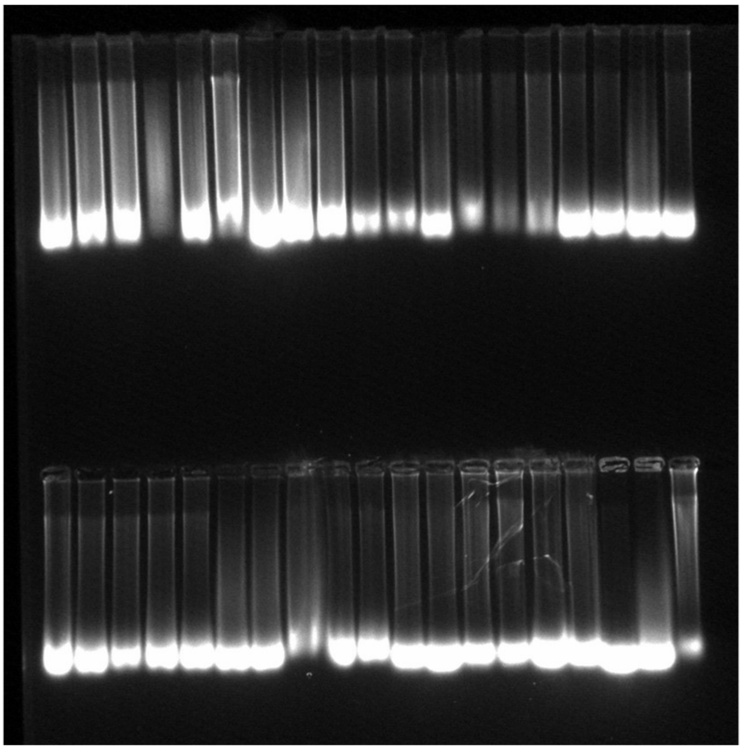

Fig. S2. Gel showing degraded total DNA extracted from randomly selected invertebrate samples.

Fig. S3. Gels showing COI1 amplification in the collected samples by universal jgLCO1490/jgHCO2198 primers. NC= negative controls. Marker: 50 bp.

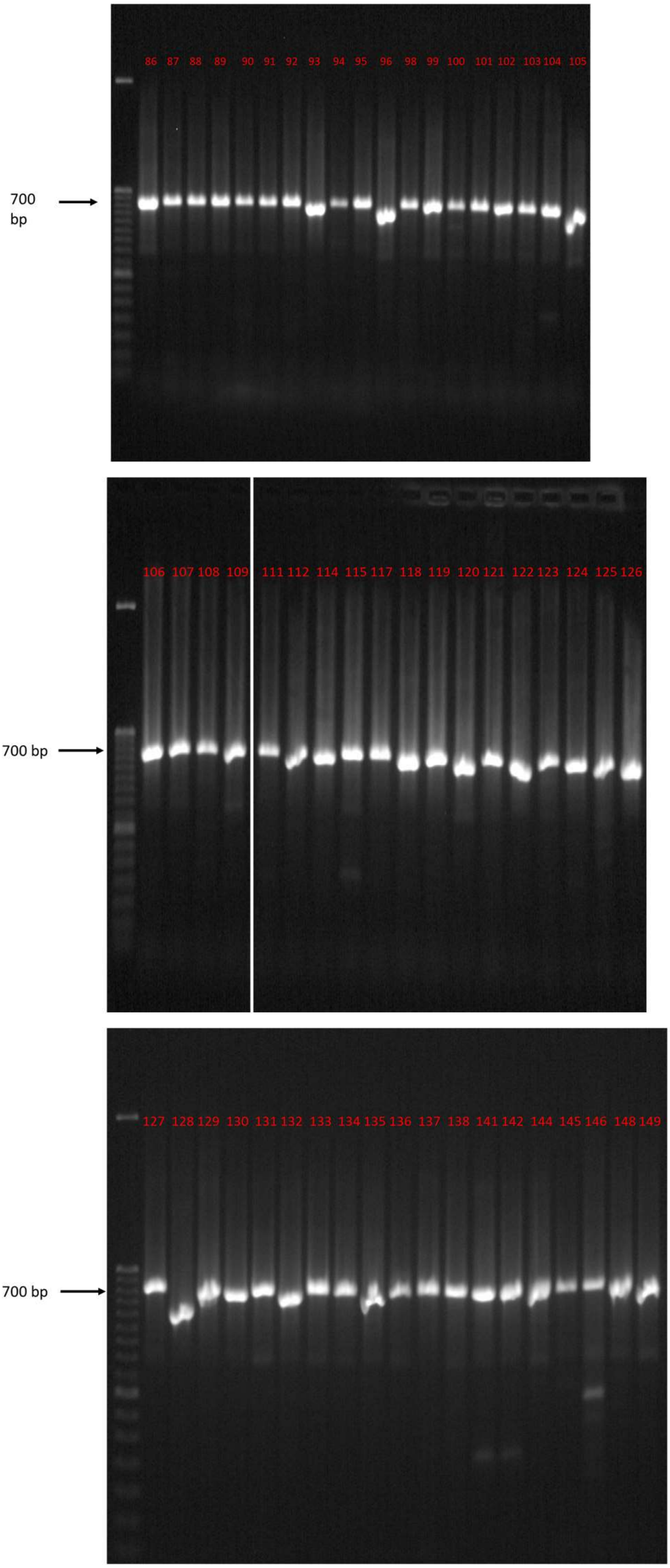

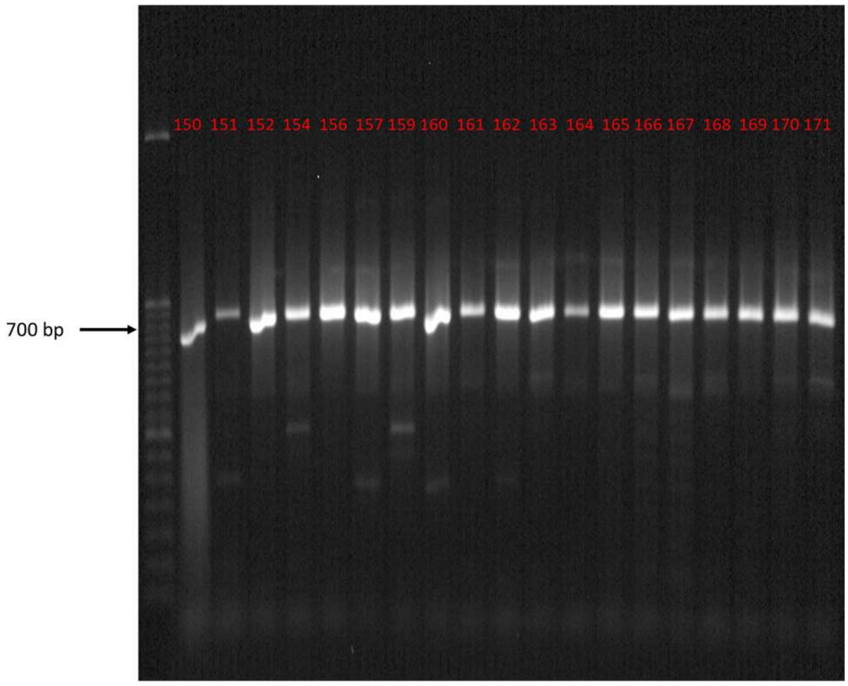
150 151 152 154 156 157 159 160 161 162 163 164 165 166 167 168 169 170 171
700 bp

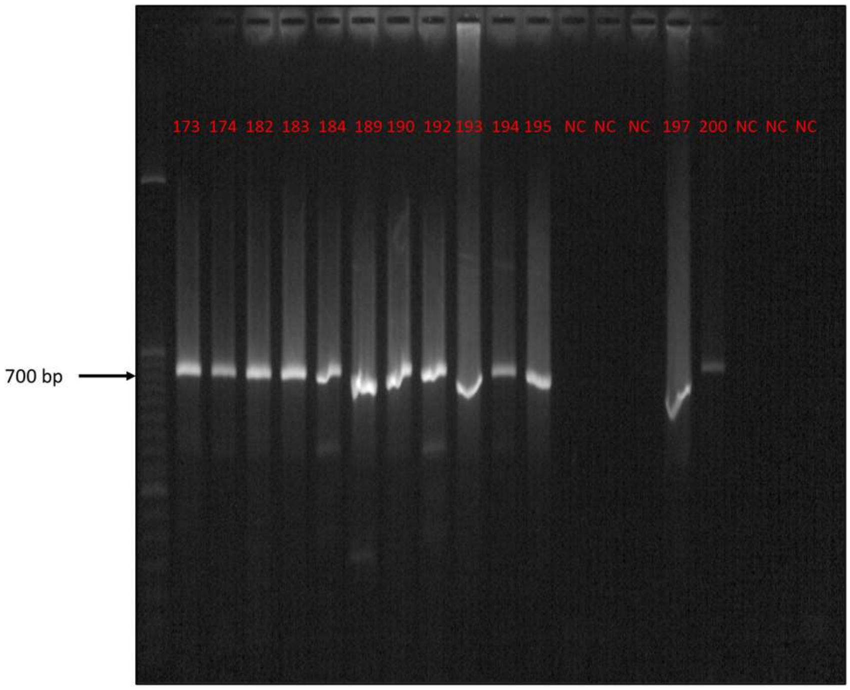
173 174 182 183 184 189 190 192 193 194 195 NC NC NC 197 200 NC NC NC
700 bp

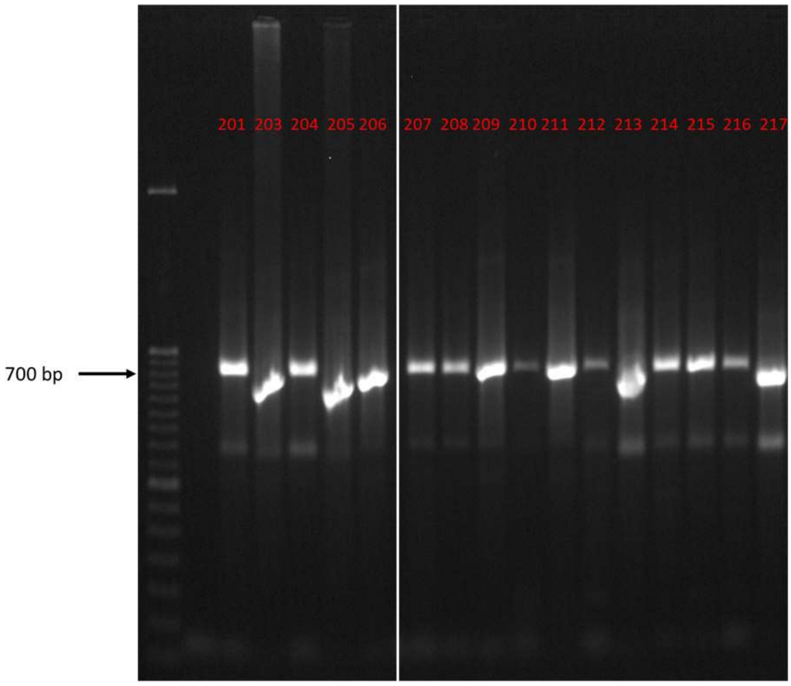
201 203 204 205 206 207 208 209 210 211 212 213 214 215 216 217
700 bp

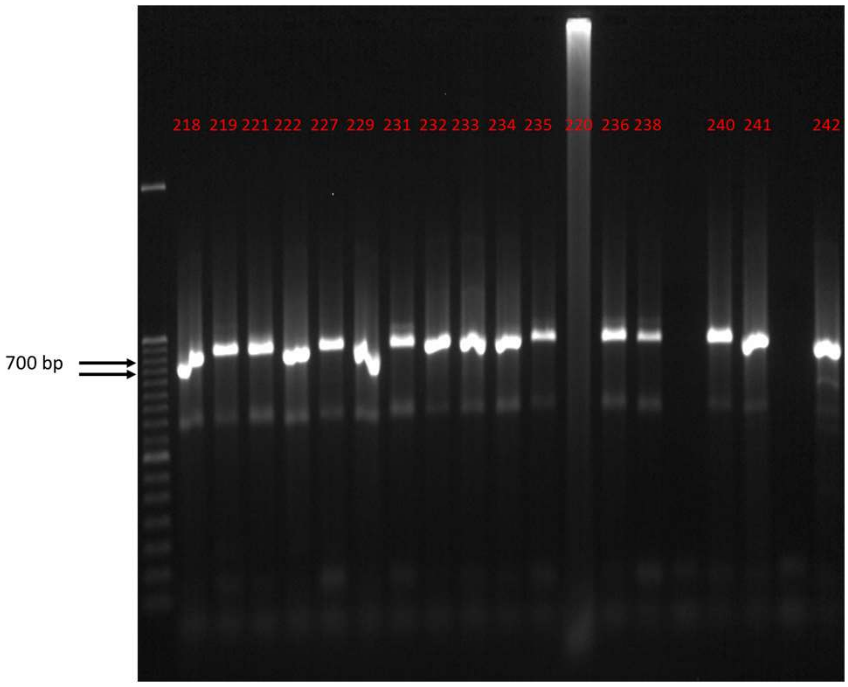
218 219 221 222 227 229 231 232 233 234 235 220 236 238 240 241 242
700 bp

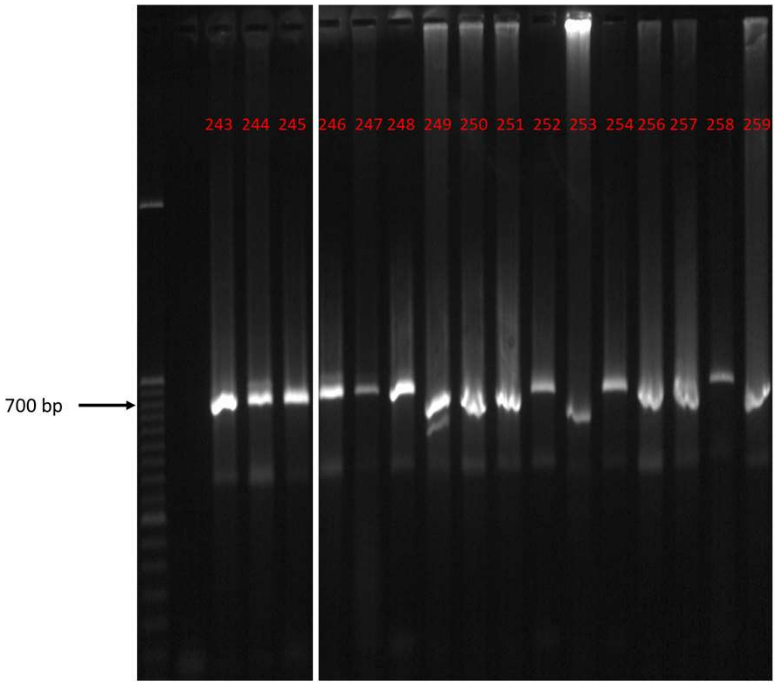
243 244 245 246 247 248 249 250 251 252 253 254 256 257 258 259
700 bp

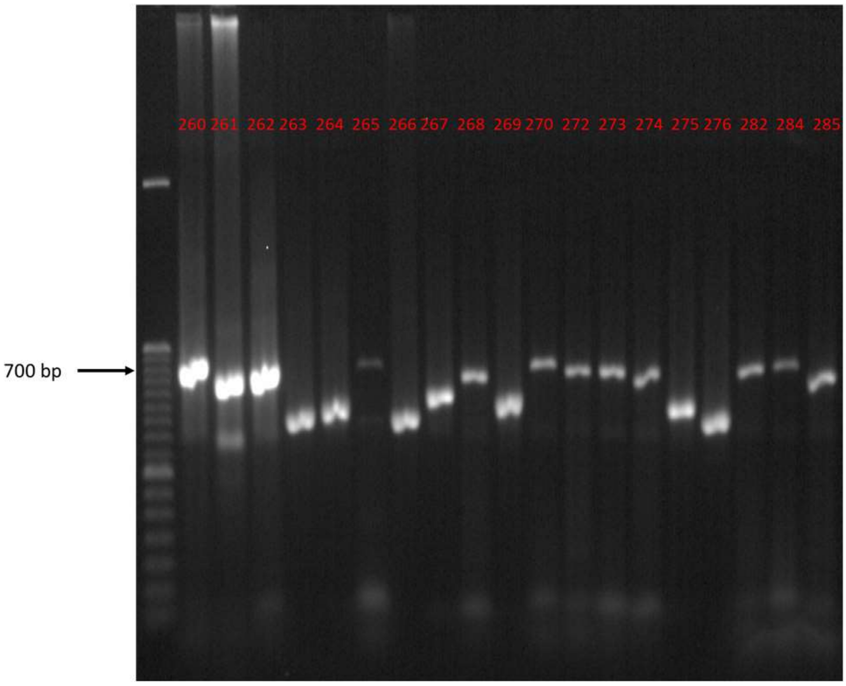
260 261 262 263 264 265 266 267 268 269 270 272 273 274 275 276 282 284 285
700 bp

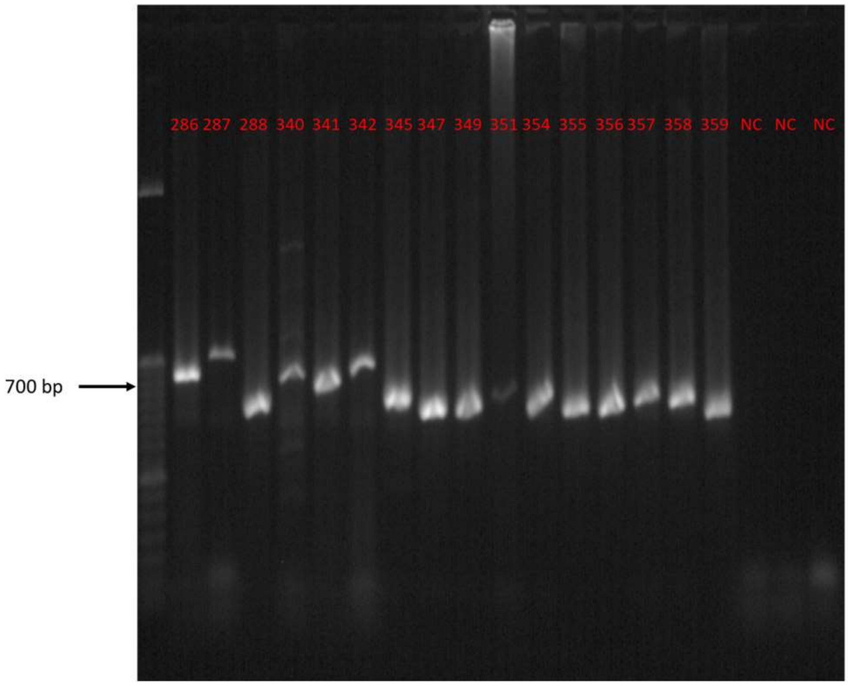
286 287 288 340 341 342 345 347 349 351 354 355 356 357 358 359 NC NC NC
700 bp

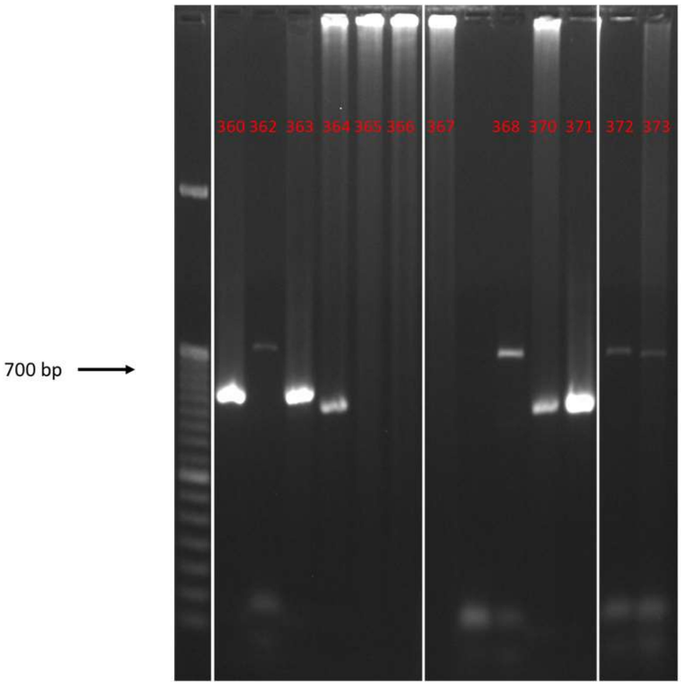
360 362 363 364 365 366 367 368 370 371 372 373
700 bp

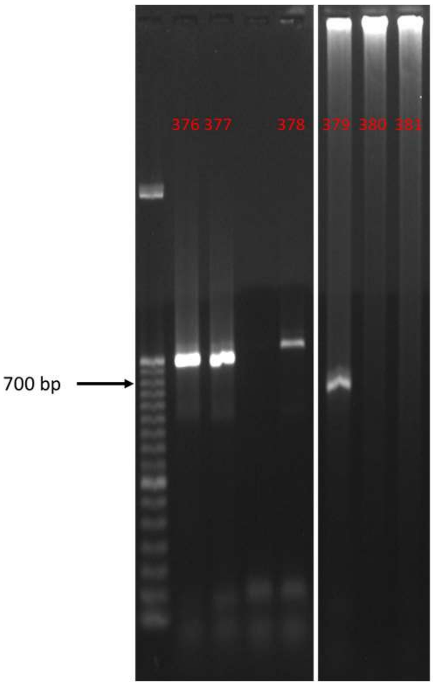
376 377 378 379 380 381
700 bp

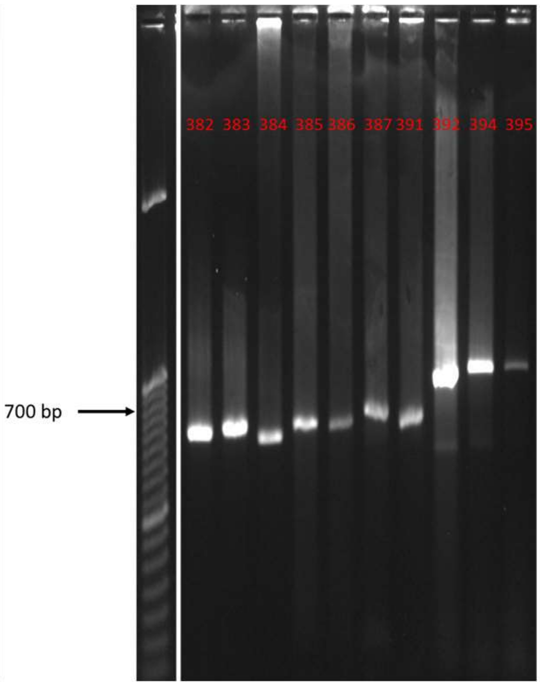
382 383 384 385 386 387 391 392 394 395
700 bp

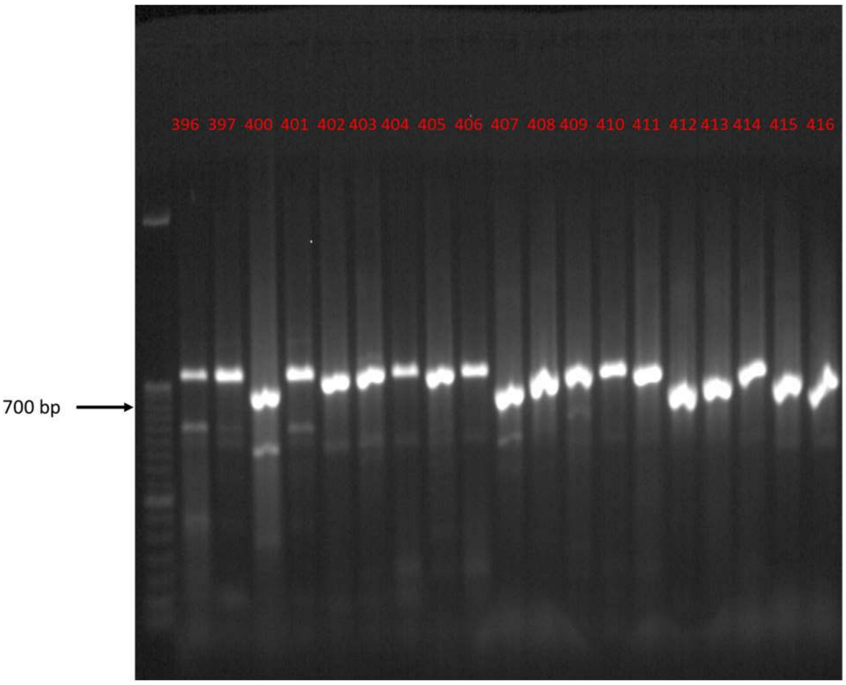
396 397 400 401 402 403 404 405 406 407 408 409 410 411 412 413 414 415 416
700 bp

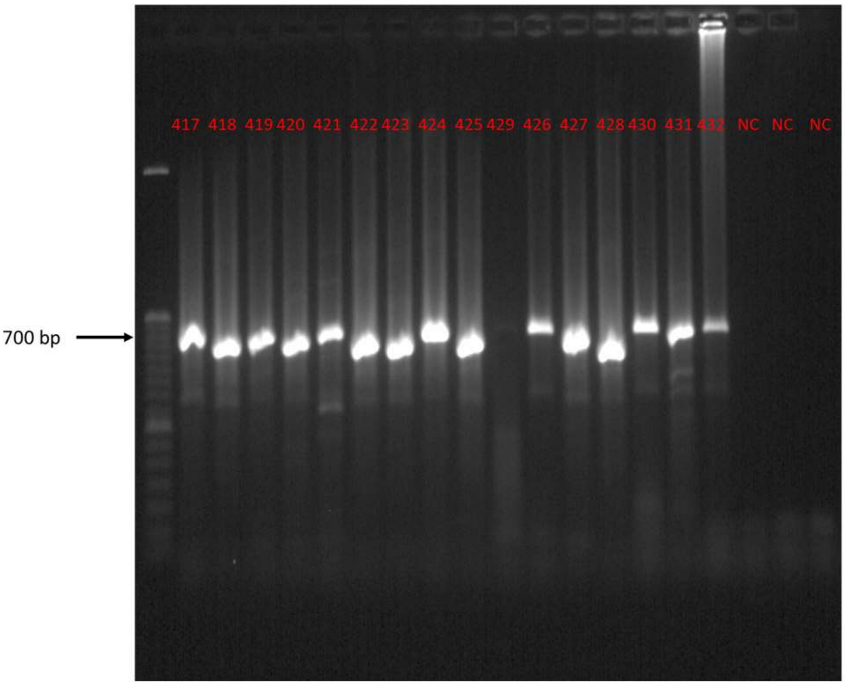
417 418 419 420 421 422 423 424 425 429 426 427 428 430 431 432 NC NC NC
700 bp

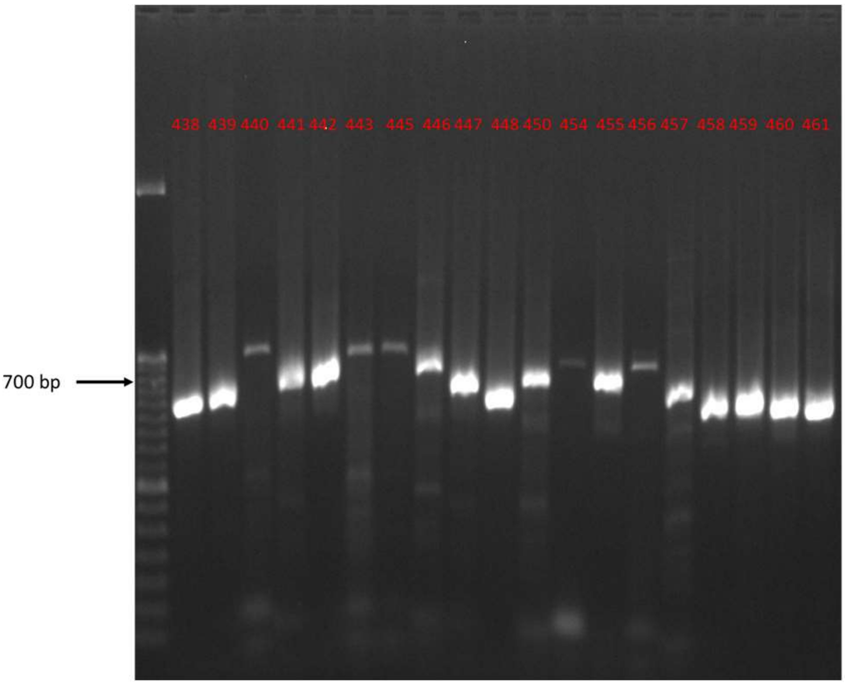
438 439 440 441 442 443 445 446 447 448 450 454 455 456 457 458 459 460 461
700 bp

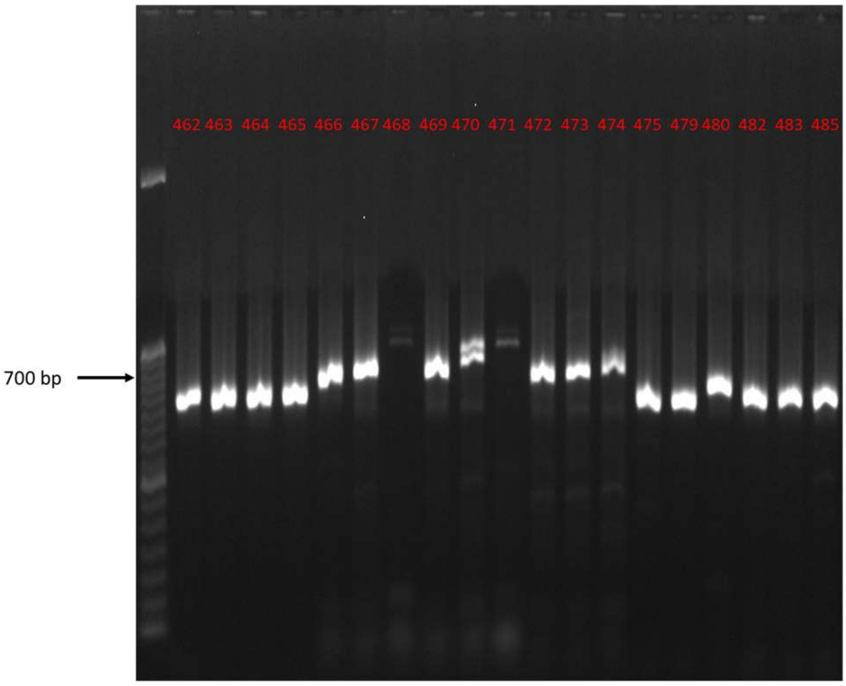
462 463 464 465 466 467 468 469 470 471 472 473 474 475 479 480 482 483 485
700 bp

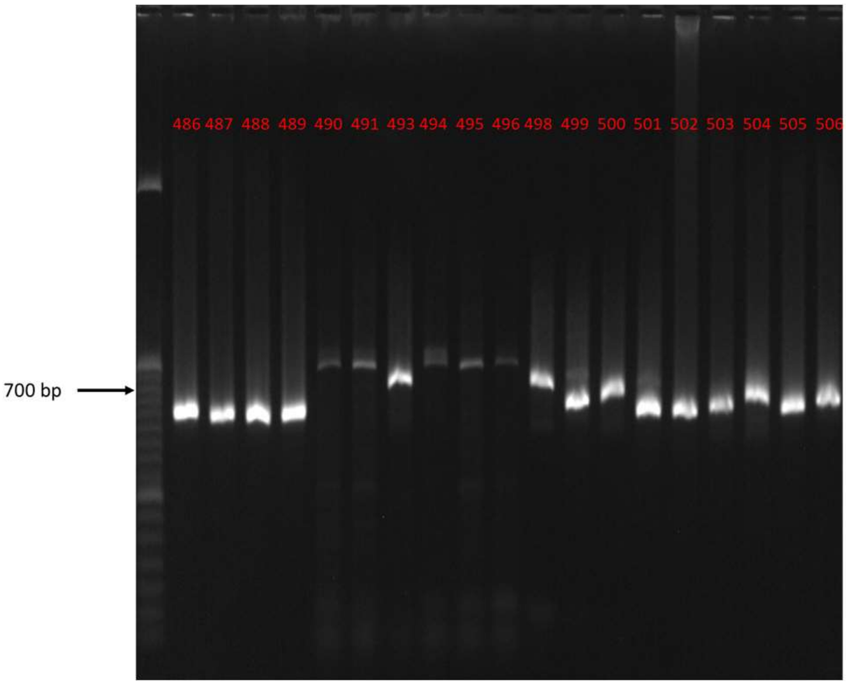
486 487 488 489 490 491 493 494 495 496 498 499 500 501 502 503 504 505 506
700 bp

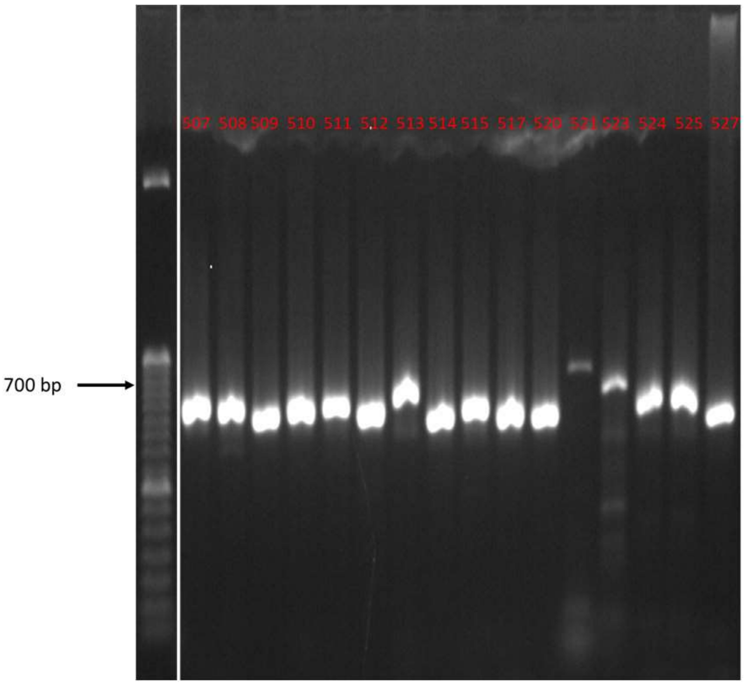
507 508 509 510 511 512 513 514 515 517 520 521 523 524 525 527
700 bp

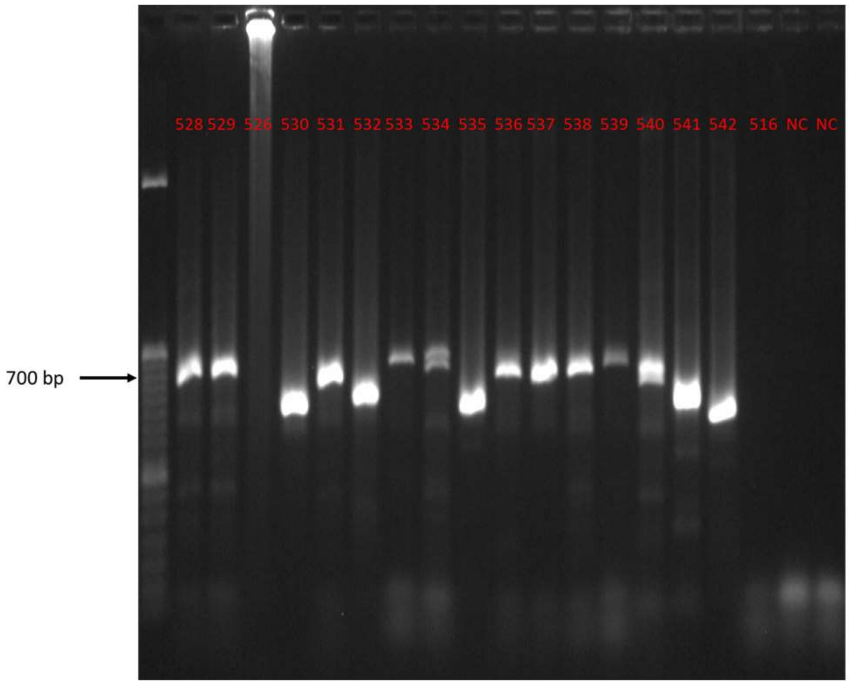
700 bp
528 529 526 530 531 532 533 534 535 536 537 538 539 540 541 542 516 NC NC

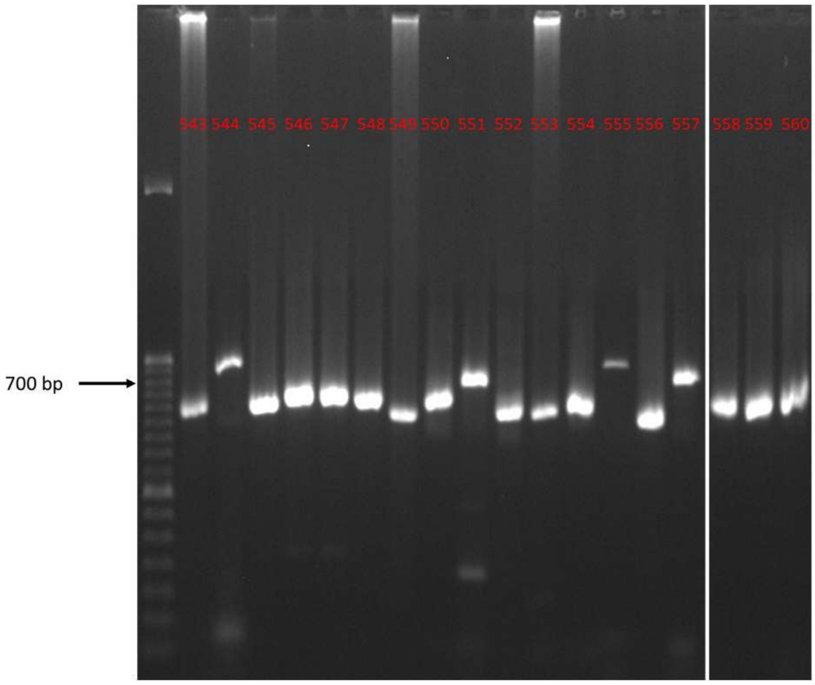
700 bp
543 544 545 546 547 548 549 550 551 552 553 554 555 556 557 558 559 560

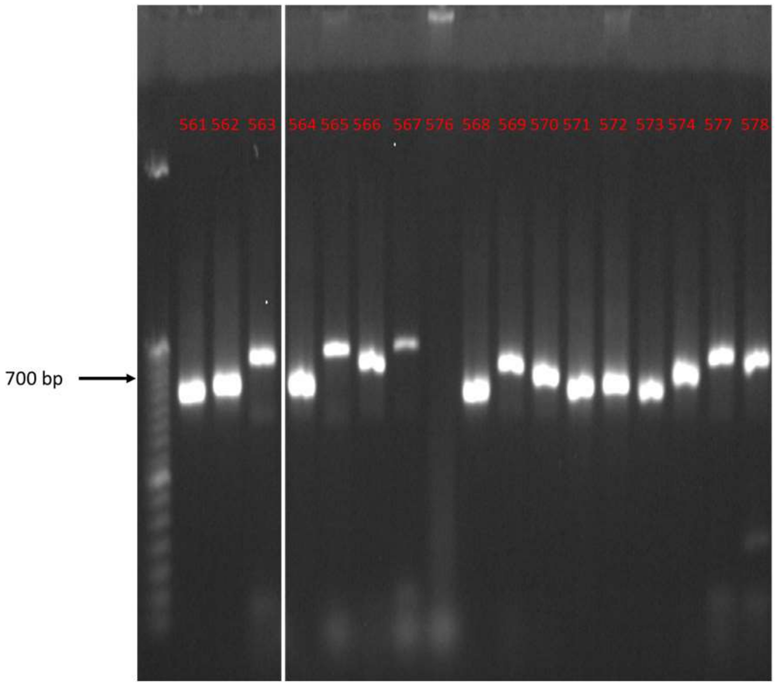
700 bp
561 562 563 564 565 566 567 576 568 569 570 571 572 573 574 577 578

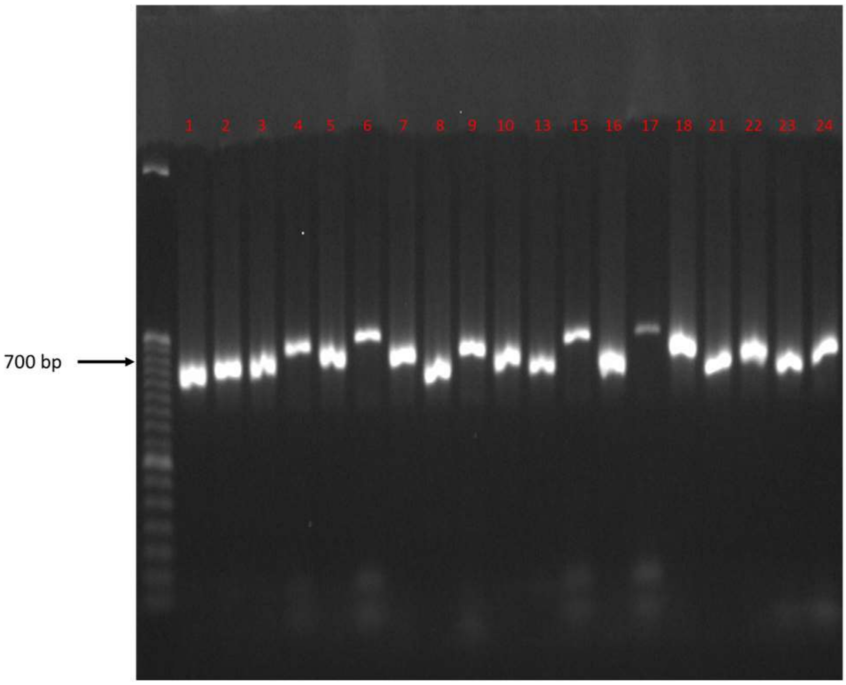
700 bp
1 2 3 4 5 6 7 8 9 10 13 15 16 17 18 21 22 23 24

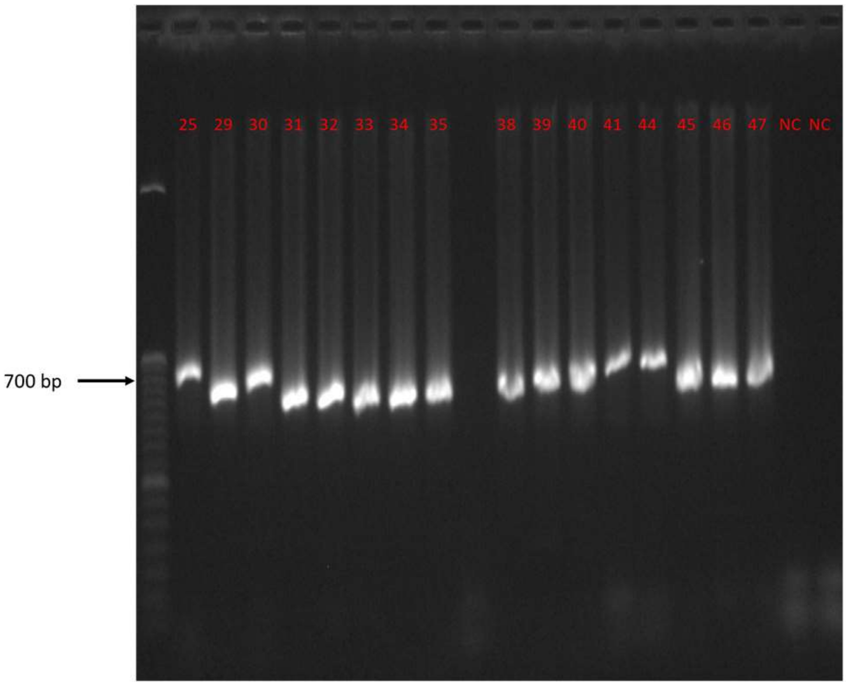
700 bp
25 29 30 31 32 33 34 35 38 39 40 41 44 45 46 47 NC NC

Fig. S4. Specificity test. Gels showing amplification of representative samples from different taxa, including positive controls *P. lividus* and *A. lixula* and negative controls (NC). Marker: 50 bp.

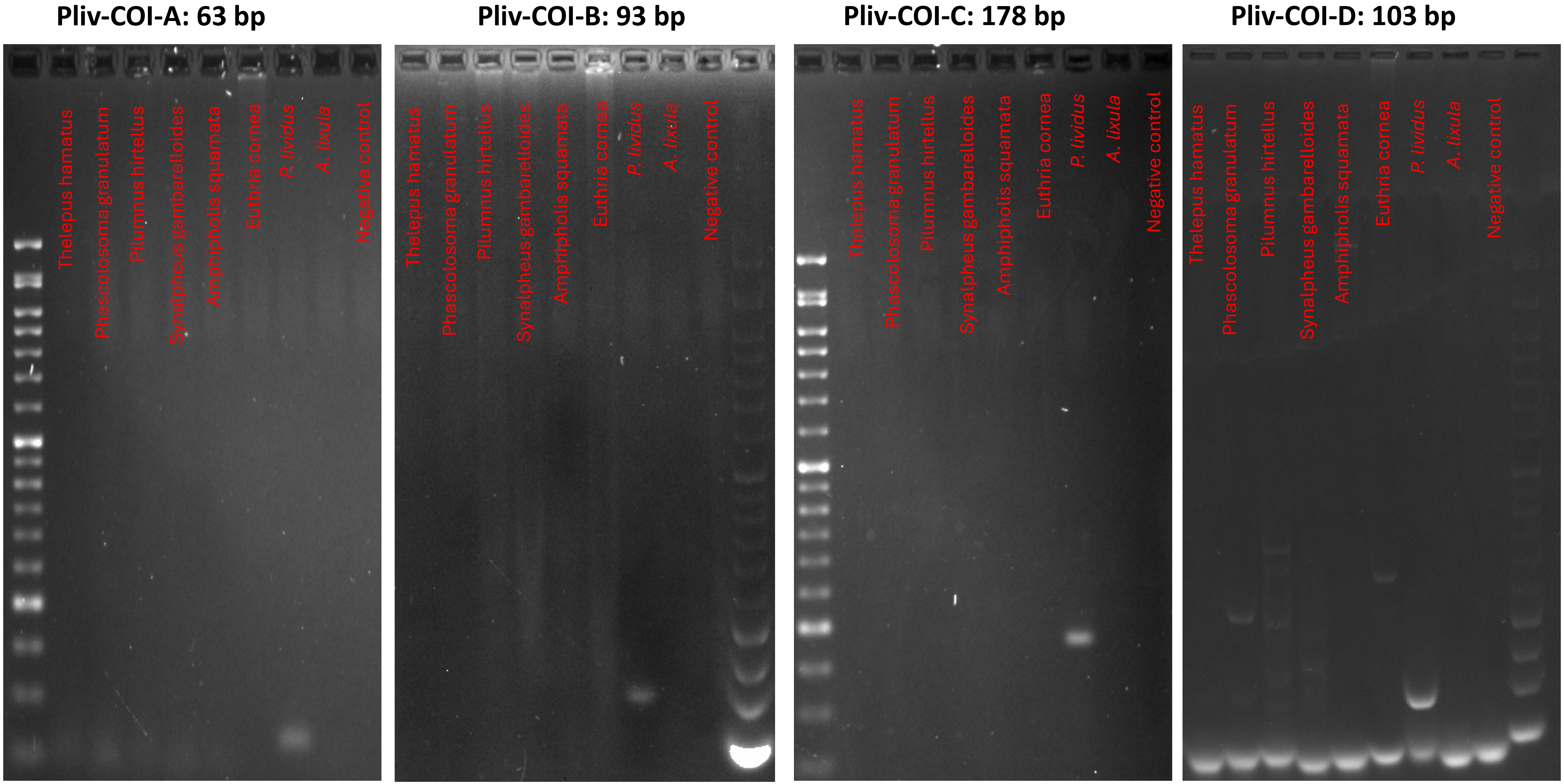

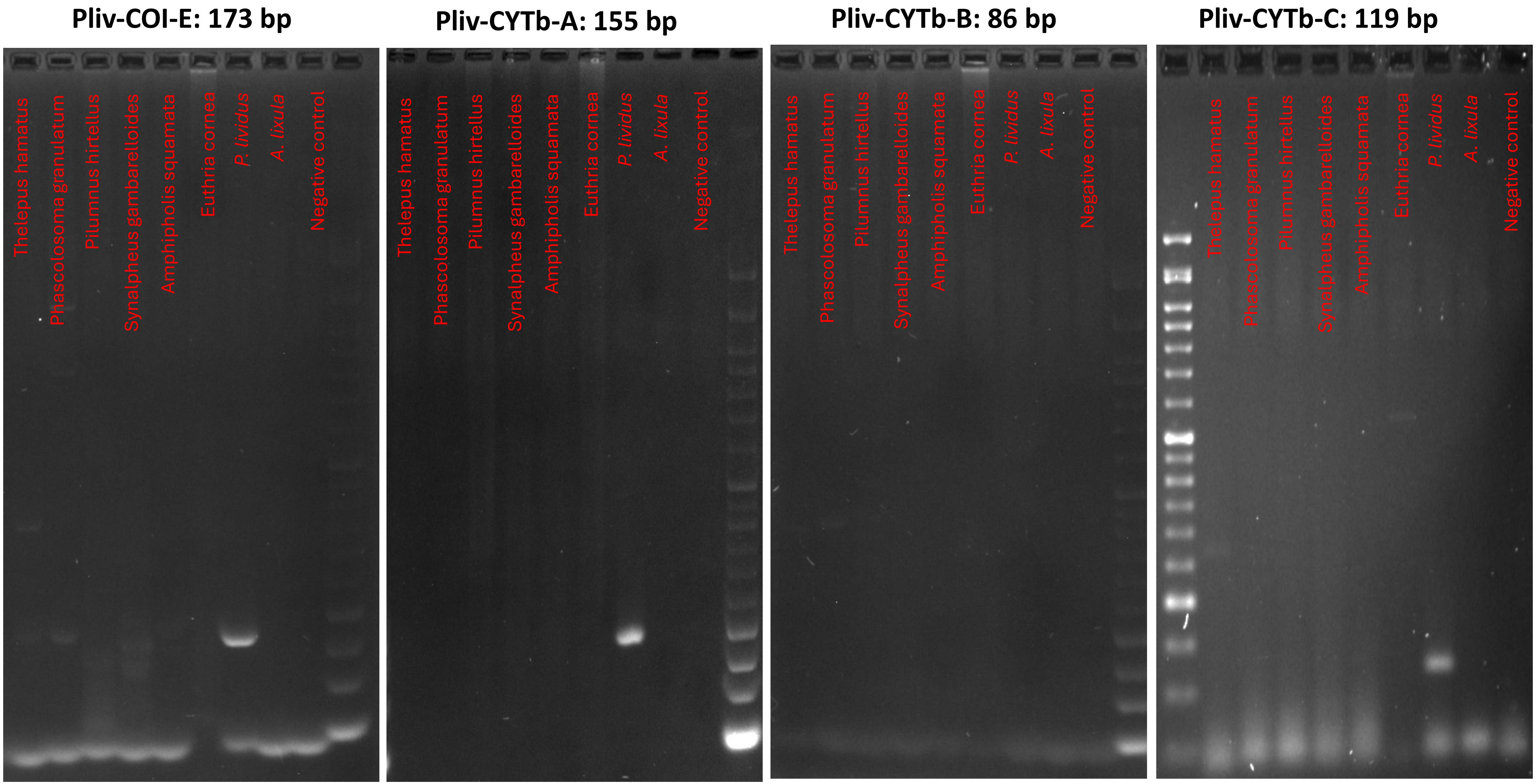
Pliv-COI-E: 173 bp
Pliv-CYTb-A: 155 bp
Pliv-CYTb-B: 86 bp
Pliv-CYTb-C: 119 bp
Thelepus hamatus
Phascolosoma granulatum
Pilumnus hirtellus
Synalpheus gambarelloides
Amphipholis squamata
Euthria cornea
P. lividus
A. lixula
Negative control

**Pliv-16S: 161 bp**

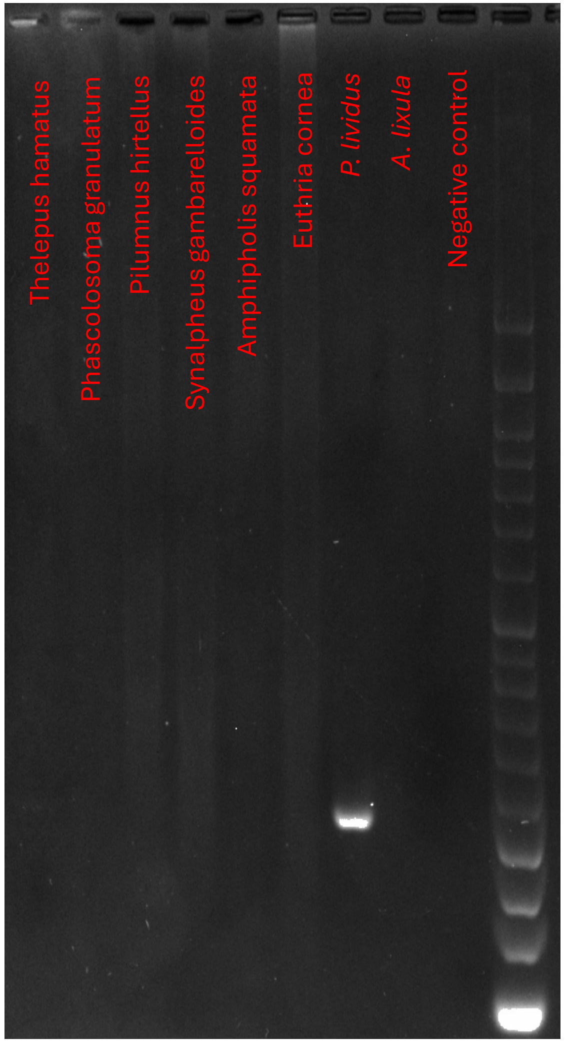

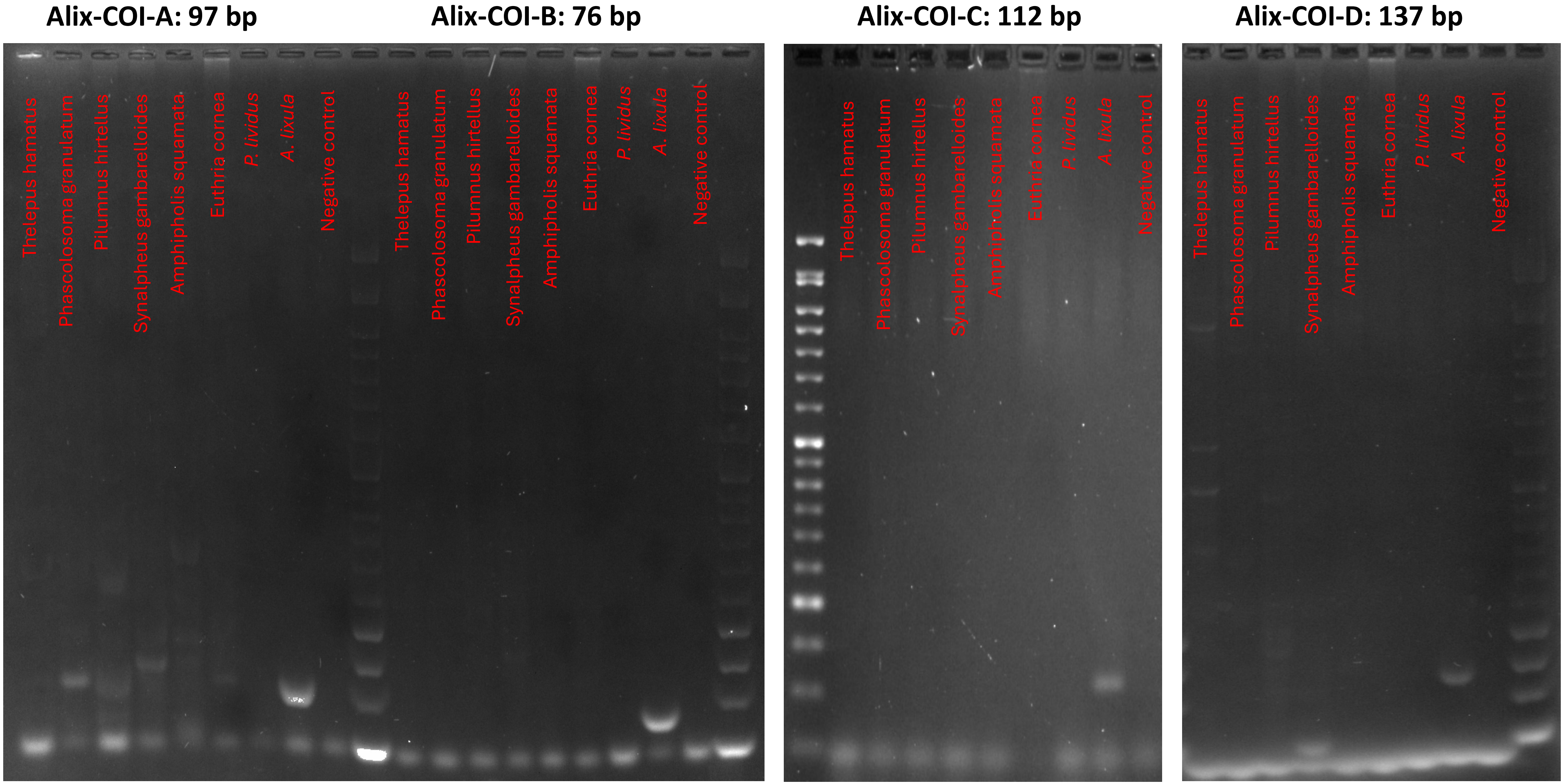

Alix-COI-A: 97 bp
Alix-COI-B: 76 bp
Alix-COI-C: 112 bp
Alix-COI-D: 137 bp
Thelepus hamatus
Phascolosoma granulatum
Pilumnus hirtellus
Synalpheus gambarelloides
Amphipholis squamata
Euthria cornea
P. lividus
A. lixula
Negative control

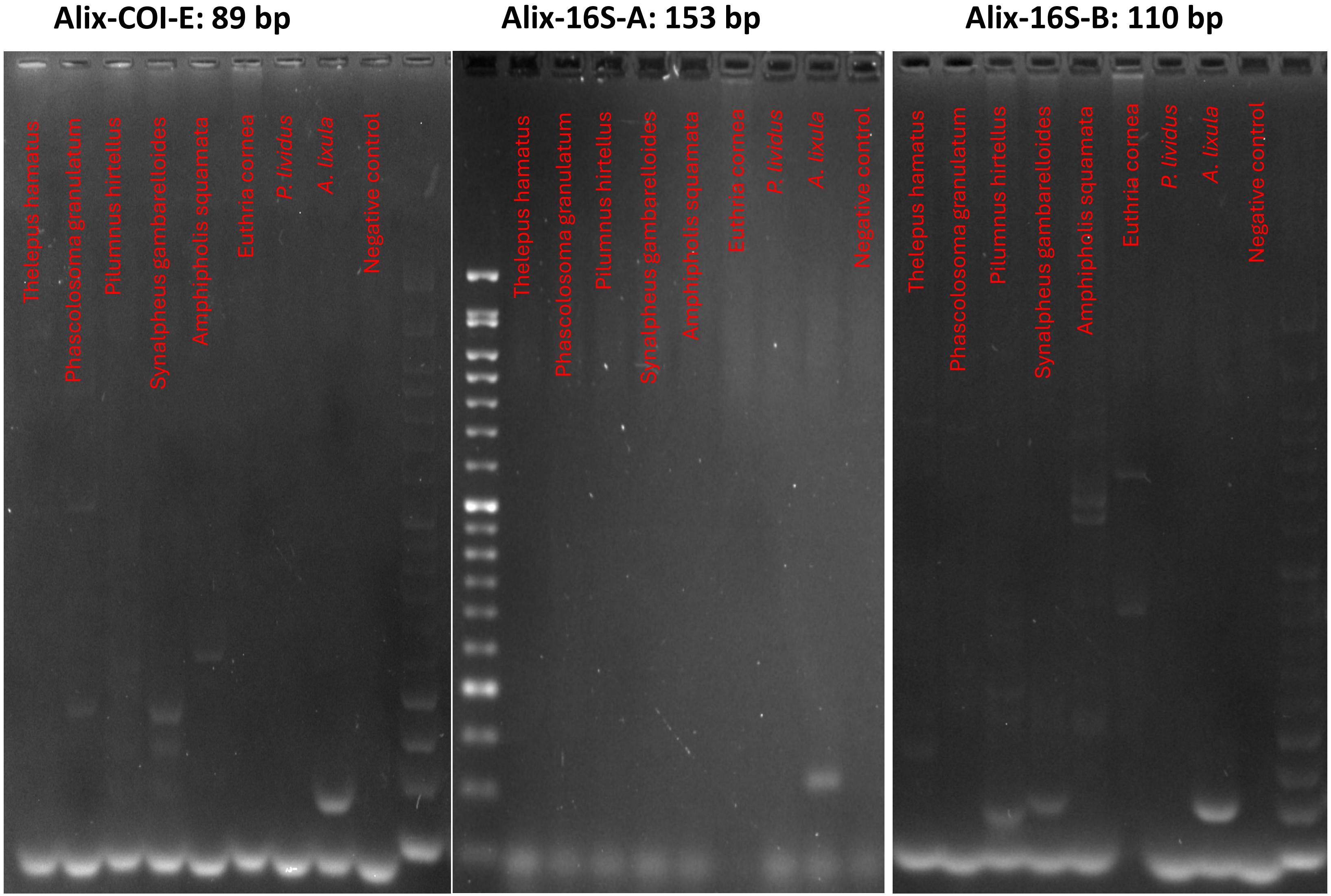
Alix-COI-E: 89 bp
Alix-16S-A: 153 bp
Alix-16S-B: 110 bp
Thelepus hamatus
Phascolosoma granulatum
Pilumnus hirtellus
Synalpheus gambarelloides
Amphipholis squamata
Euthria cornea
P. lividus
A. lixula
Negative control

Fig. S5. Sensitivity test. Gels showing amplification of *P. lividus* and *A. lixula* DNA in a dilution concentration series. NC= negative controls. Marker: 50 bp.

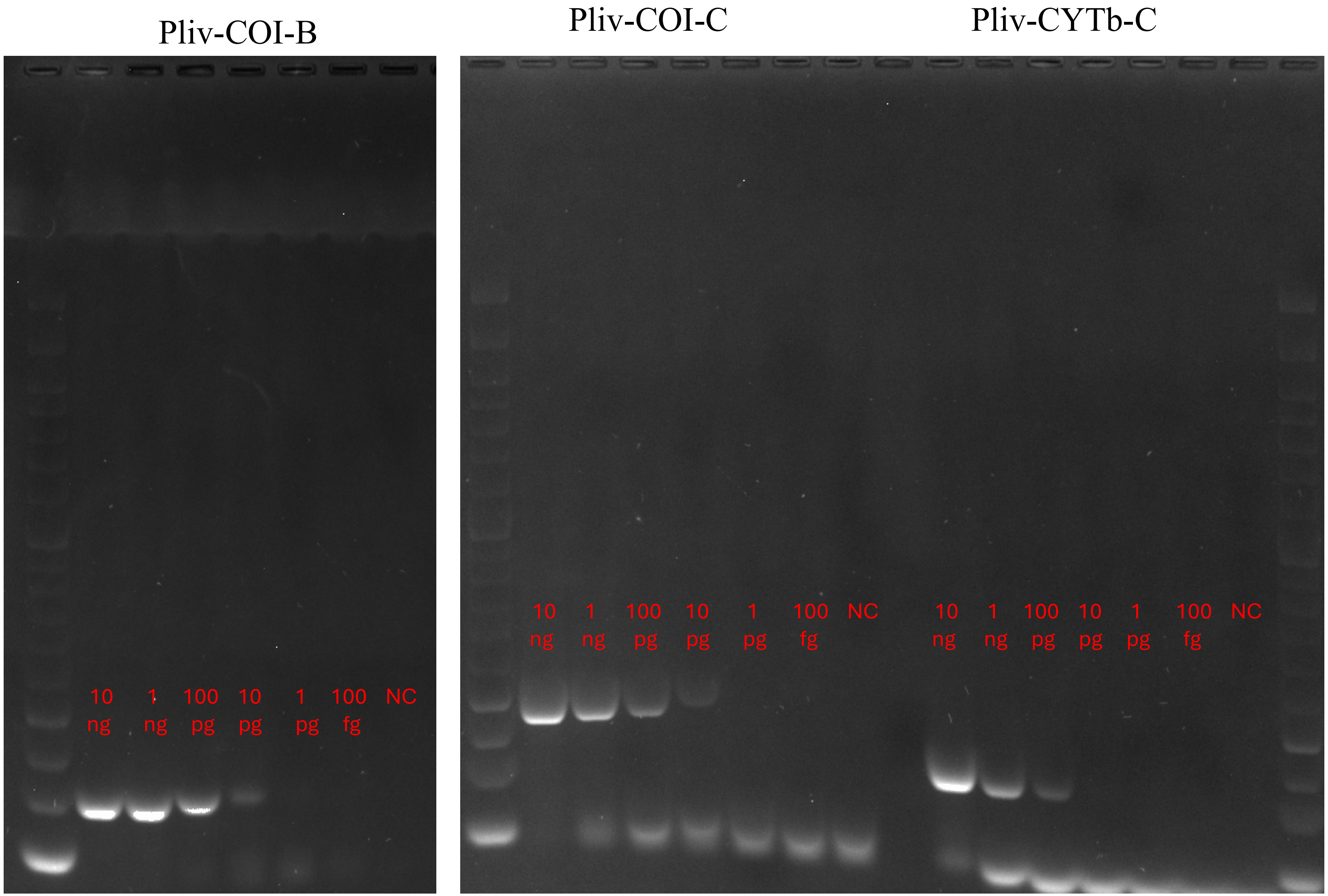

Pliv-16S Pliv-CYTb-A

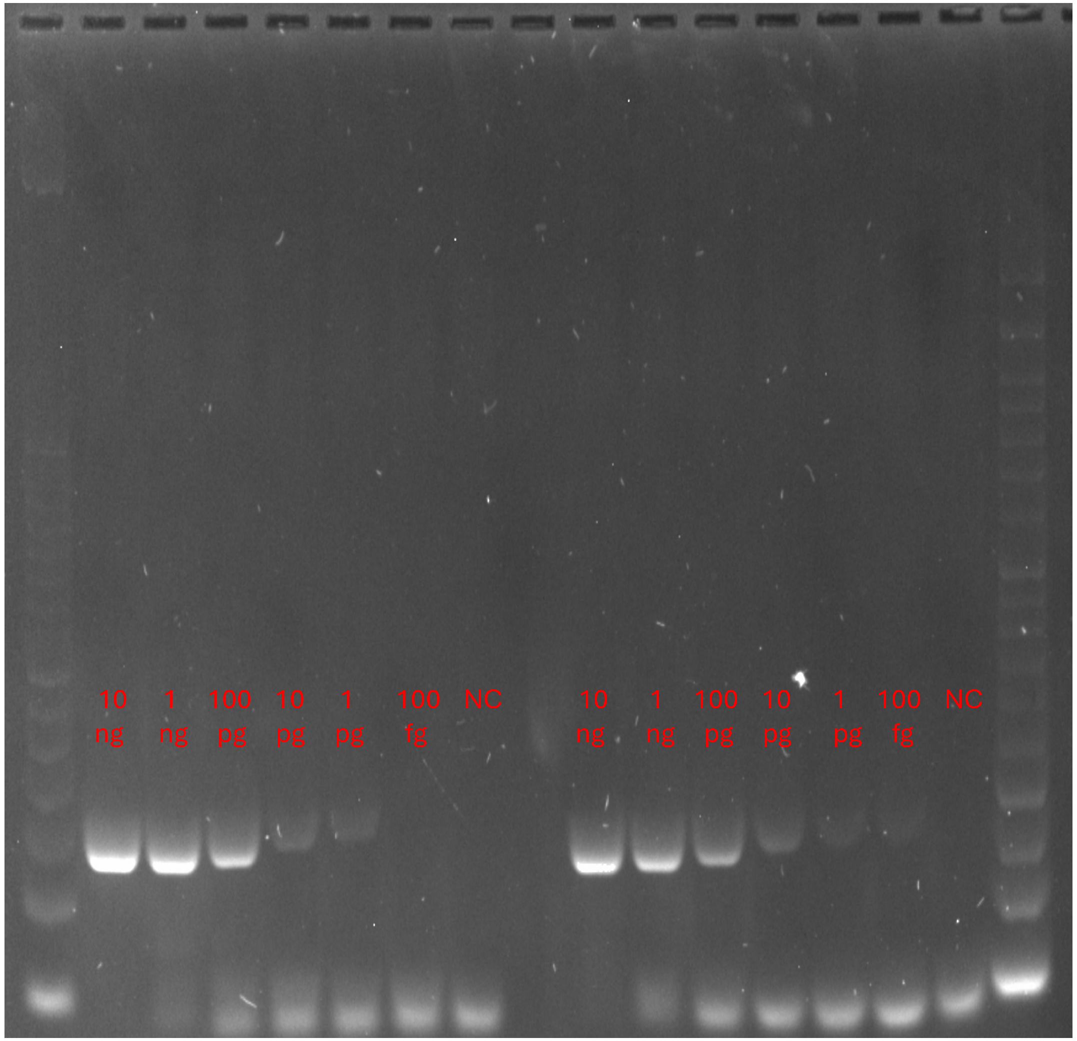

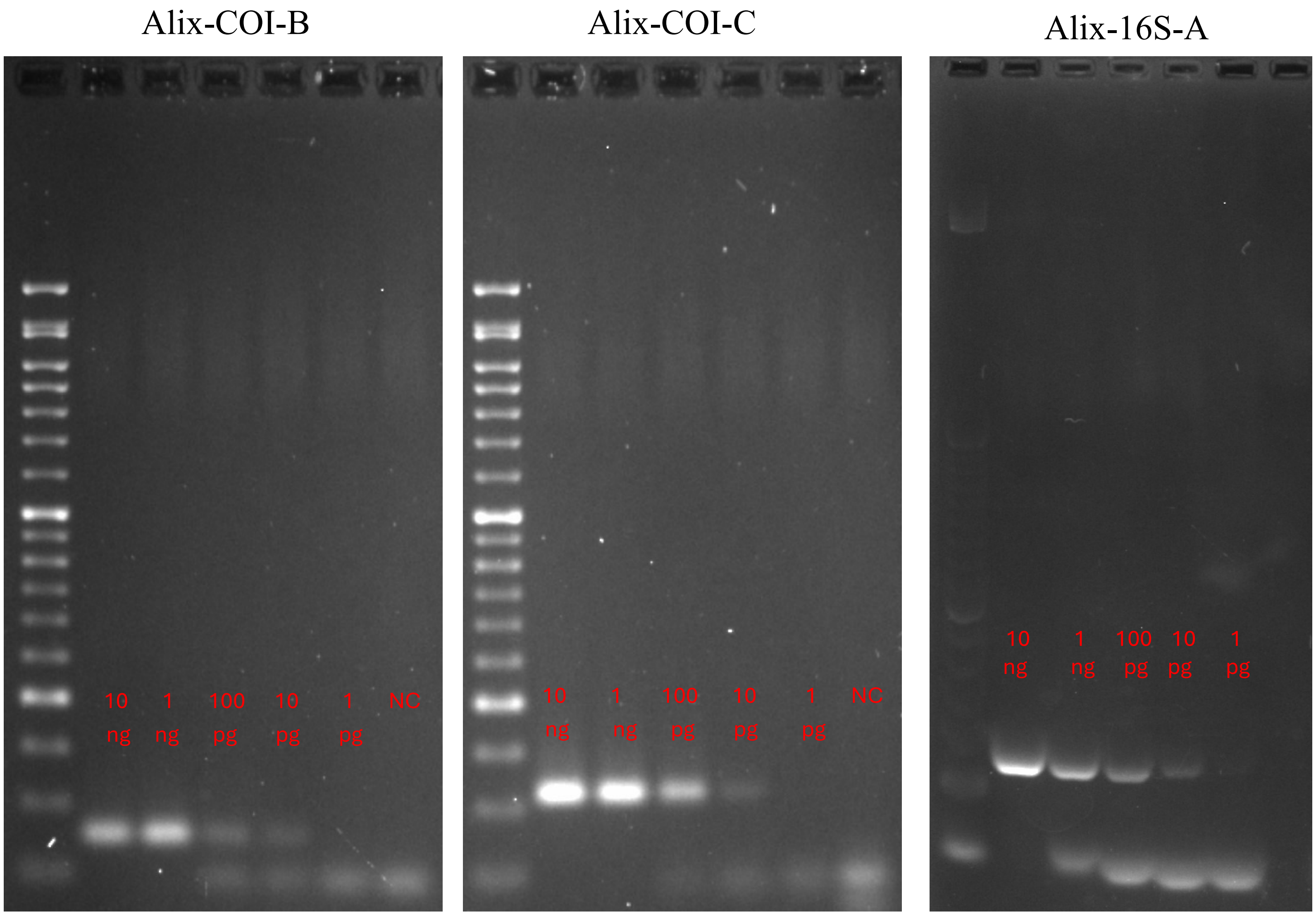
Alix-COI-B
10 ng
1 ng
100 pg
10 pg
1 pg
NC
Alix-COI-C
10 ng
1 ng
100 pg
10 pg
1 pg
NC
Alix-16S-A
10 ng
1 ng
100 pg
10 pg
1 pg

Fig. S6. Inhibition test. Gels showing amplification of *P. lividus* and *A. lixula* DNA when mixed with 1,000 thousand-fold excess of predator's DNA. NC= negative controls. Marker: 50 bp.

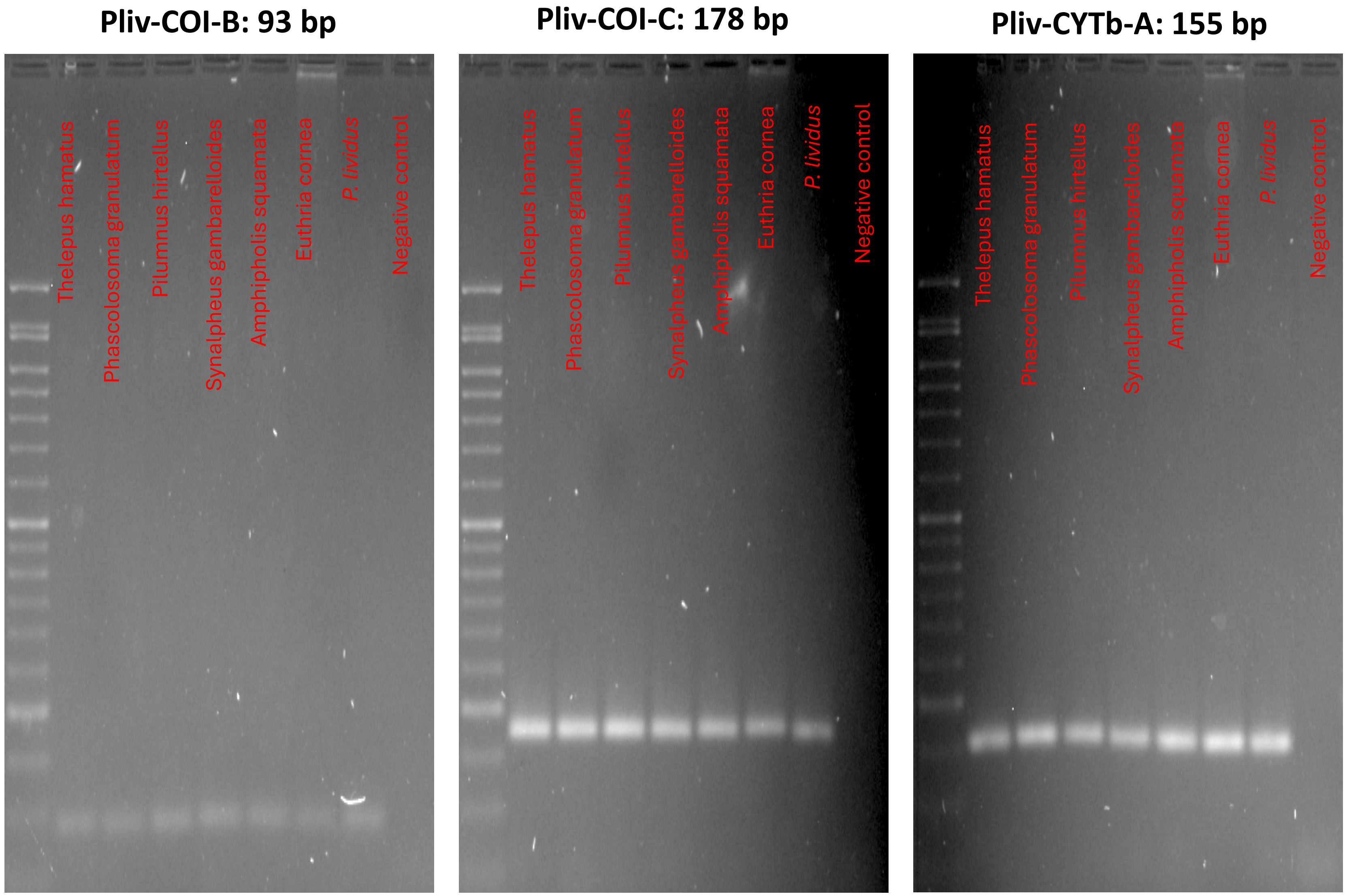

**Pliv-CYTb-C: 119 bp**

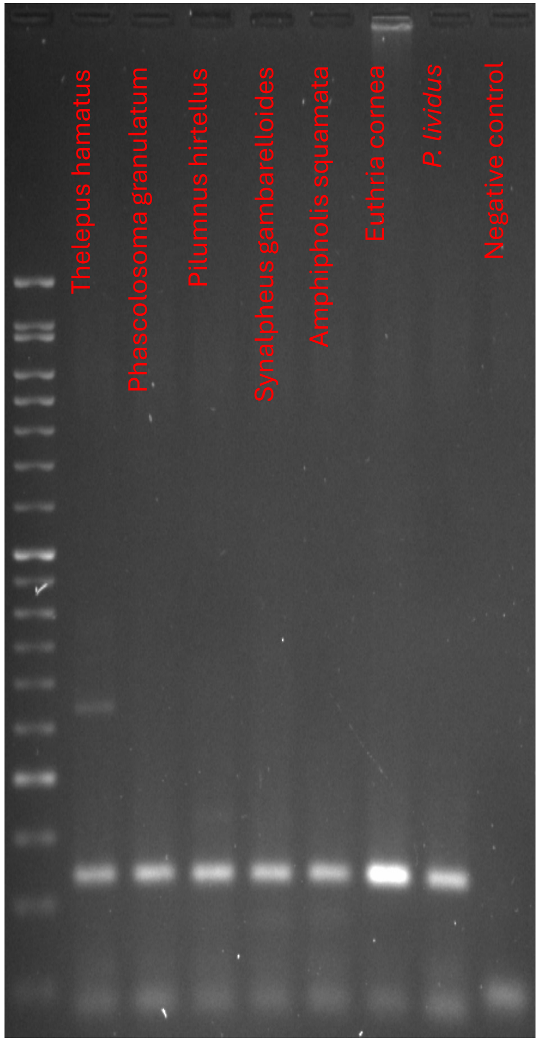


**Pliv-16S: 161 bp**

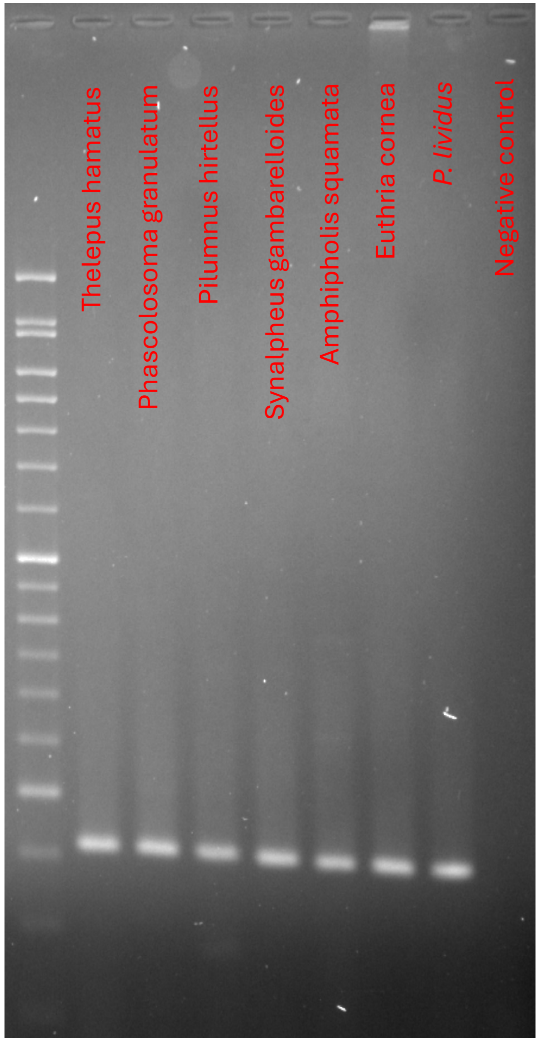

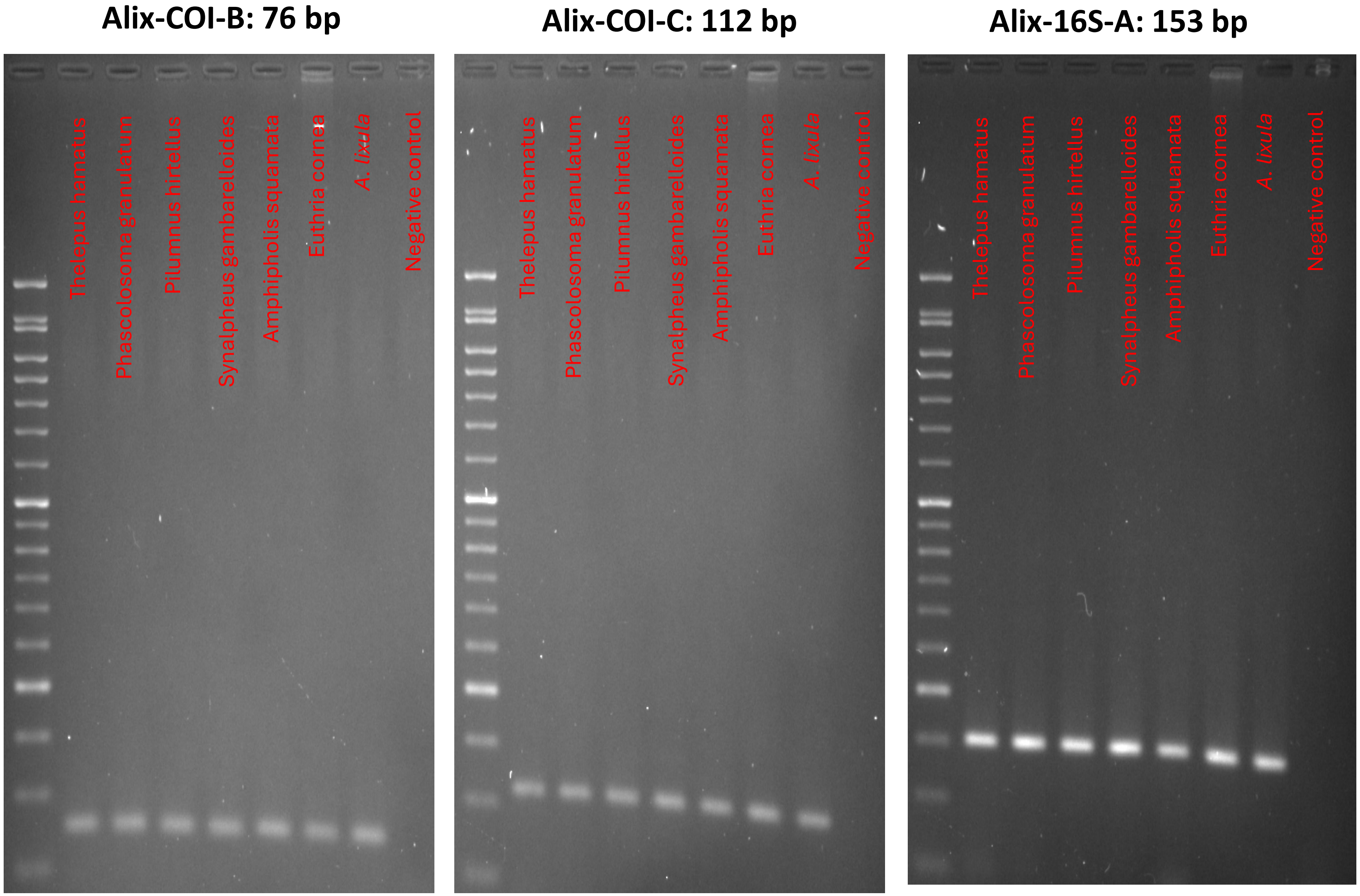
Alix-COI-B: 76 bp
Alix-COI-C: 112 bp
Alix-16S-A: 153 bp
Thelepus hamatus
Phascolosoma granulatum
Pilumnus hirtellus
Synalpheus gambarelloides
Amphipholis squamata
Euthria cornea
A. lixula
Negative control

Fig. S7. Gels showing amplification from urchin settlers' DNA with *P. lividus* and *A. lixula* specific primers. NC= negative controls. Marker: 50 bp.

### Pliv-16S

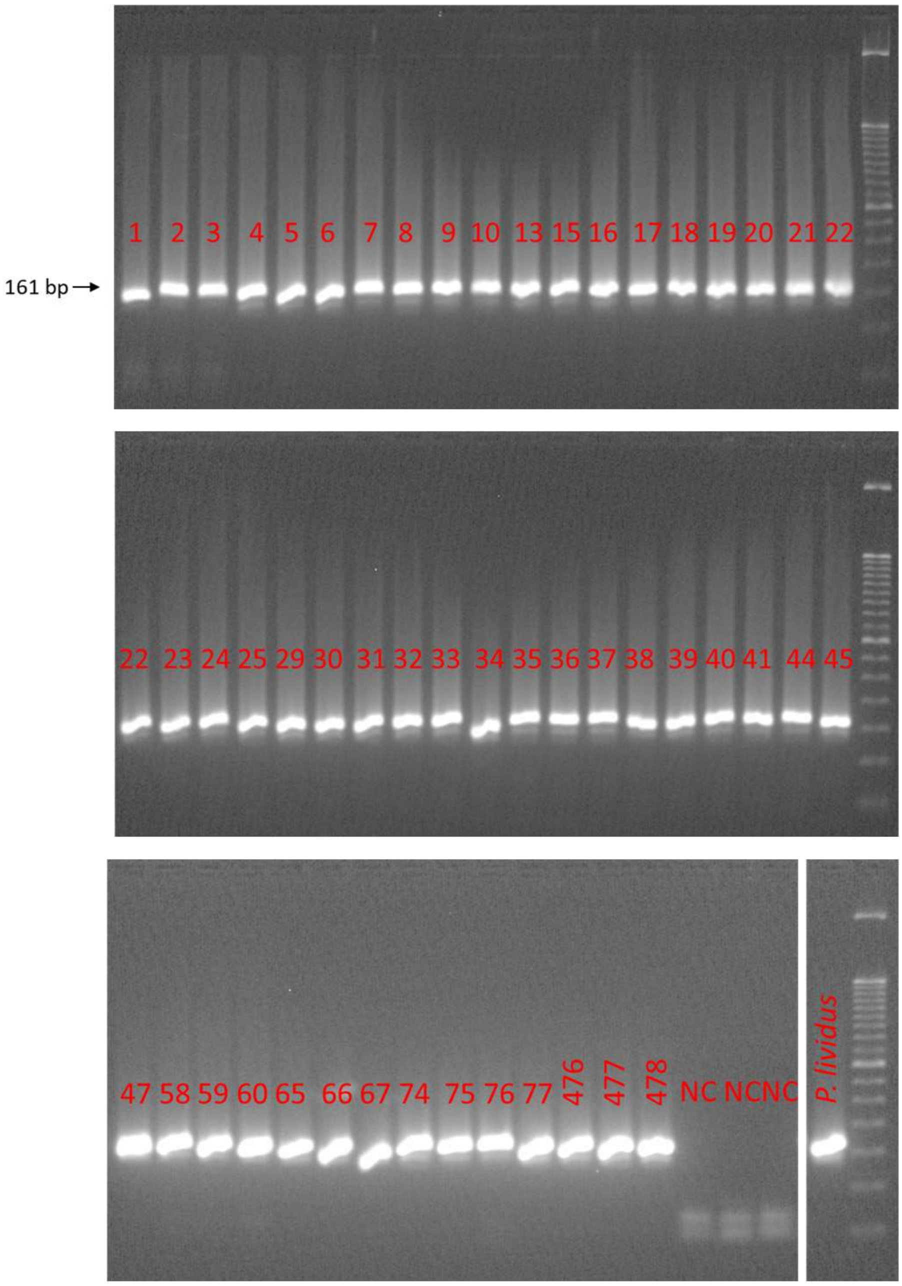

Pliv-CYTb-A

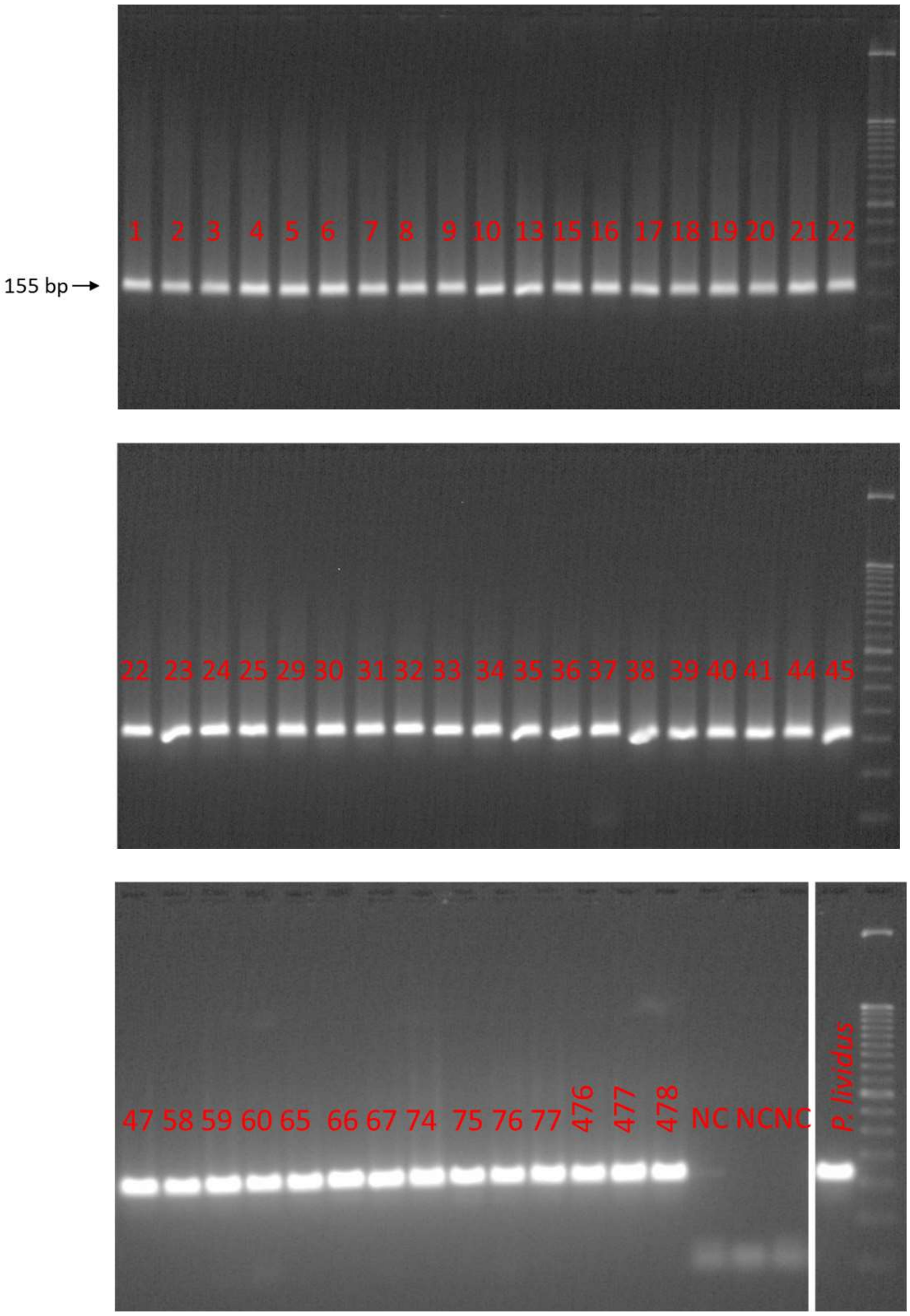

155 bp
1 2 3 4 5 6 7 8 9 10 13 15 16 17 18 19 20 21 22
22 23 24 25 29 30 31 32 33 34 35 36 37 38 39 40 41 44 45
47 58 59 60 65 66 67 74 75 76 77 476 477 478 NC NC NC P. lividus

Pliv-COI-B

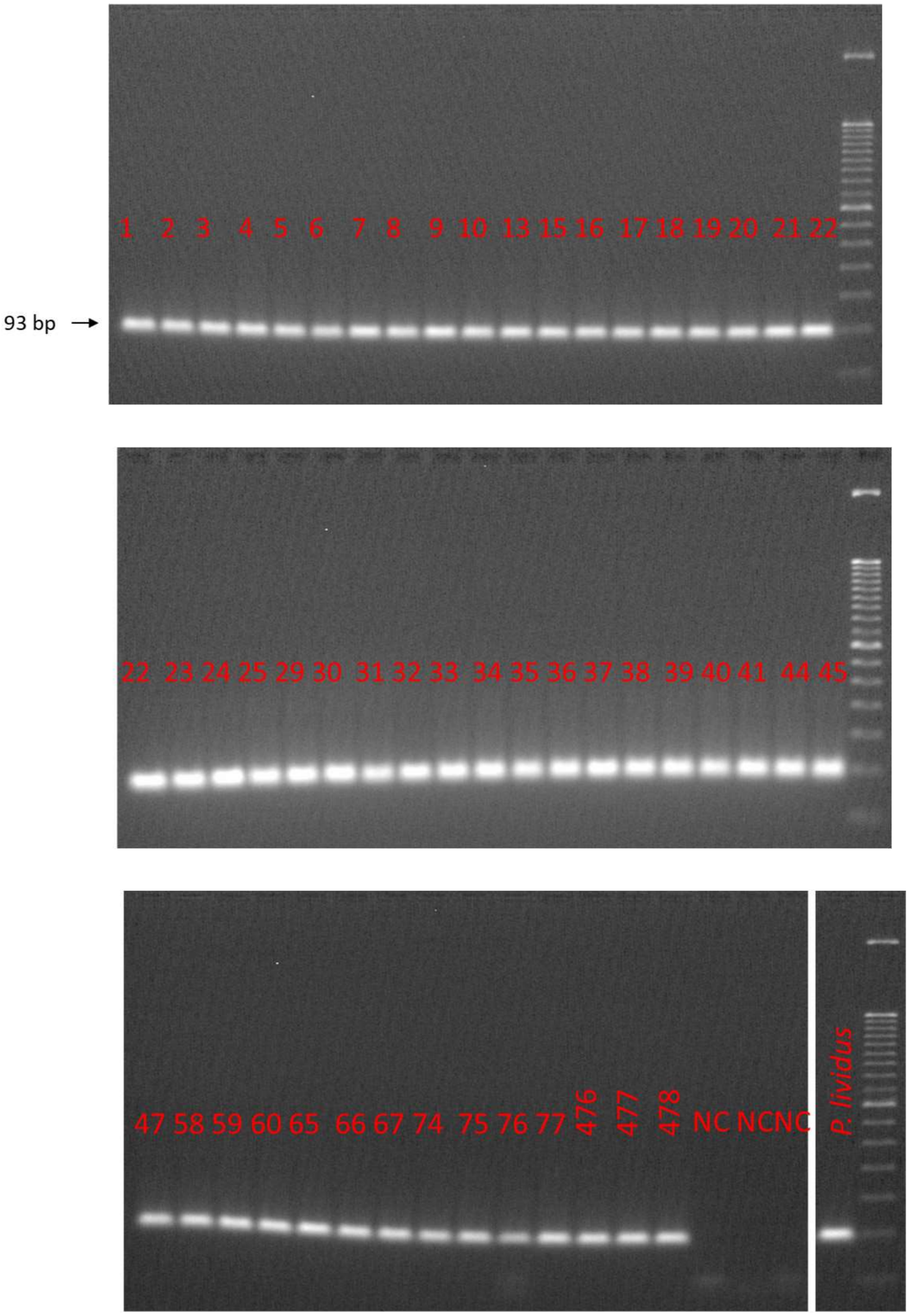
93 bp
1 2 3 4 5 6 7 8 9 10 13 15 16 17 18 19 20 21 22
22 23 24 25 29 30 31 32 33 34 35 36 37 38 39 40 41 44 45
47 58 59 60 65 66 67 74 75 76 77 476 477 478 NC NC NC
*P. lividus*

Al-16S- A

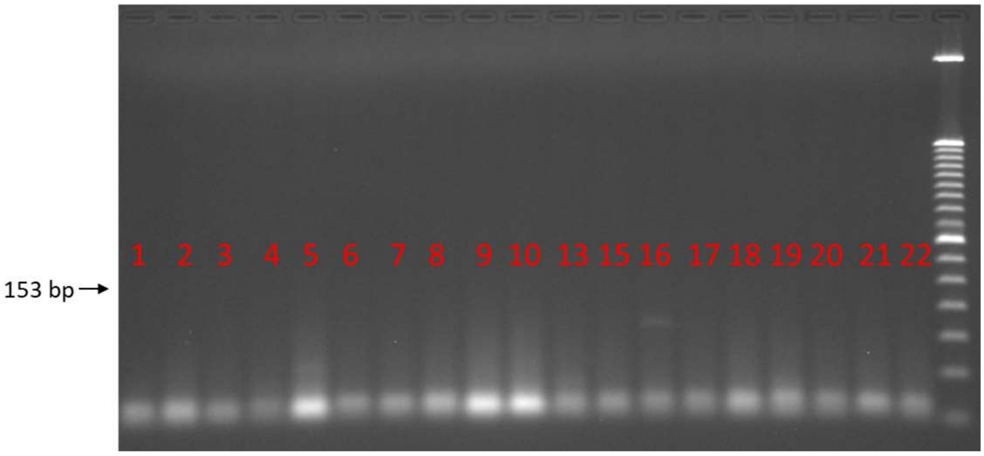
153 bp
1 2 3 4 5 6 7 8 9 10 13 15 16 17 18 19 20 21 22

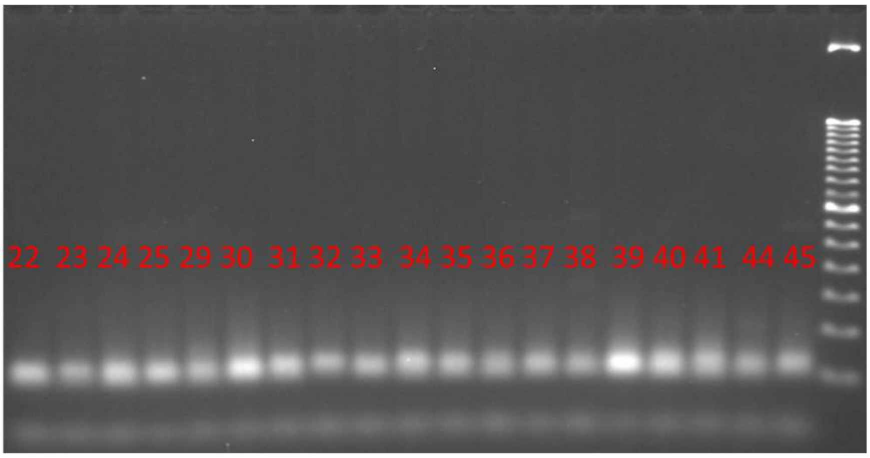
22 23 24 25 29 30 31 32 33 34 35 36 37 38 39 40 41 44 45

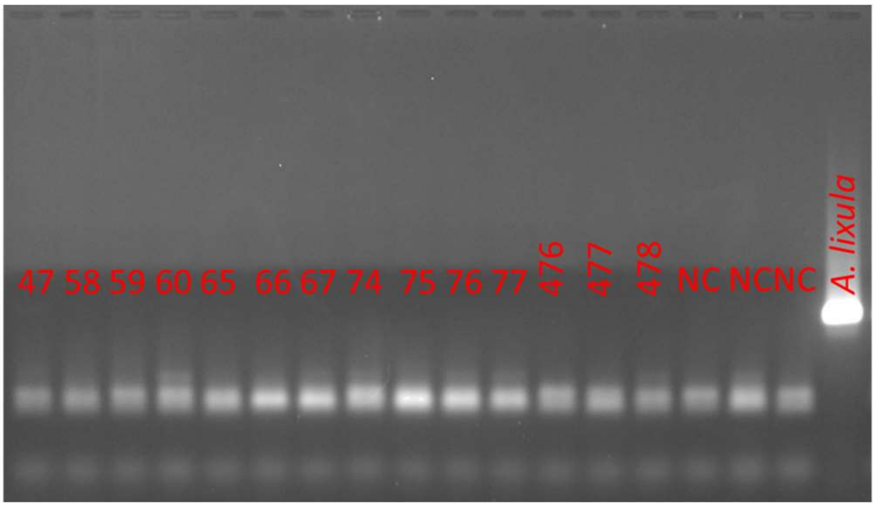
47 58 59 60 65 66 67 74 75 76 77 476 477 478 NC NC NC *A. lixula*
